# Structural basis for the inhibition of PRC2 by active transcription histone posttranslational modifications

**DOI:** 10.1101/2024.02.09.579730

**Authors:** Trinity Cookis, Alexandria Lydecker, Paul Sauer, Vignesh Kasinath, Eva Nogales

## Abstract

Polycomb repressive complex 2 (PRC2) is an epigenetic regulator essential for embryonic development and maintenance of cell identity that trimethylates histone H3 at lysine 27 (H3K27me3) leading to gene silencing. PRC2 is regulated by association with protein cofactors and crosstalk with histone posttranslational modifications. Trimethylated histone H3 K4 (H3K4me3) and K36 (H3K36me3) localize to sites of active transcription where H3K27me3 is absent and inhibit PRC2 activity through unknown mechanisms. Using cryo-electron microscopy we reveal that histone H3 tails modified with H3K36me3 engage poorly with the PRC2 active site and preclude its effective interaction with chromatin, while the H3K4me3 modification binds to the allosteric site in the EED subunit, acting as an antagonist that competes with allosteric activators required for the spreading of the H3K27me3 repressive mark. Thus, the location along the H3 tail of the H3K4me3 and H3K36me3 modifications allow them to target two essential requirements for efficient trimethylation of histone H3K27. We further show that the JARID2 cofactor modulates PRC2 activity in the presence of these histone modifications.

## Introduction

The dynamic recognition, deposition, and removal of posttranslational modifications (PTMs) on histone proteins facilitate the establishment of specific gene expression programs. Many chromatin-modifying enzymes are large protein complexes with catalytic and regulatory regions capable of sensing the chromatin environment. Preexisting chromatin marks may act as recruitment platforms and/or directly stimulate or restrict the catalytic activities of chromatin-modifying enzymes. The precise recruitment, activation, and interplay between the chromatin-modifying machinery and the chromatin state are vital to defining active or repressed gene states during development and maintaining them throughout the organism’s lifespan.

Polycomb Repressive Complex 2 (PRC2) is an essential epigenetic regulator that marks genes for repression through its deposition of trimethylation on histone H3 lysine 27 (H3K27me3)^1^. It contains four core subunits (EZH1/2, EED, RBAP46/48, and SUZ12) and additionally associates with accessory proteins that impact the recruitment and catalytic functions of the complex: AEBP2 and JARID2 in PRC2.2 and EPOP/PALI1 and PHF1/MTF2/PHF19 in PRC2.1^2,3^. The interaction of PRC2 with substrate nucleosomes is mainly driven through two structural elements, the bridge helix and the CXC domain, within the catalytic subunit EZH2. These elements interact with nucleosomal DNA via electrostatic interactions and contribute to extensive contacts that help guide the histone H3 tail into the active site^4,5^.

PRC2 activity is regulated on chromatin through crosstalk with several histone PTMs. A central mechanism of PRC2 activation is the recognition of PRC2-trimethylated lysine peptides by its EED subunit, resulting in conformational changes within PRC2 that lead to the allosteric activation of the complex^4,6,7^. As a consequence, H3K27me3, the product of PRC2 enzymatic activity on nucleosomes, binds to EED and activates EZH2, therefore enabling the spreading of this modification across chromatin to establish and maintain repressed domains^4,7–10^. PRC2 also trimethylates its accessory proteins JARID2 (at K116) and PALI1 (at K1241), which can then mimic H3K27me3 to serve as allosteric activators of the complex ^6,11–14^. Recognition of allosteric activators at the EED regulatory site results in the folding of the stimulatory response motif (SRM) helix, which interacts with the catalytic SET domain of EZH2, enhancing its enzymatic activity ^4,7,10^. PRC2.2 accessory proteins, JARID2 and AEBP2, additionally mediate interactions with chromatin previously modified by PRC1 with H2AK119Ub to further promote PRC2 activity at defined genomic locations^5,15,16^.

Despite the accumulation of molecular details depicting interactions between PRC2 and chromatin in states that promote catalytic activity^5,17–19^, there is still a poor understanding of the mechanisms proposed to exclude PRC2 activity from actively transcribed chromatin, such as RNA^20–24^ and histone post-translational modifications^25,26^. The H3K4me3 and H3K36me3 histone modifications localize at promoters^27–31^ and gene bodies^28,30,32,33^ of actively transcribed genes, respectively. Genomic locations harboring these modifications are practically devoid of H3K27me3^34–36^, and in vitro biochemical assays have demonstrated that both H3K4me3 and H3K36me3 directly inhibit PRC2 enzymatic activity^5,18,25,26^. It has also been shown that JARID2 can enhance the activity of PRC2 in the presence of both H3K4me3 and H3K36me3 chromatin modifications^5,18,37^. Deciphering the crosstalk between PRC2, its accessory proteins, and the epigenetic landscape is required to understand how the activities of polycomb proteins are regulated to control gene repression and establish heterochromatin boundaries.

Here, we have used cryo-electron microscopy (cryo-EM) to determine structures of PRC2.2 (from here on PRC2) complexes engaged with nucleosome substrates containing H3K4me3 or H3K36me3, both in the presence and absence of the methylated JARID2 K116. We discovered that H3K4me3- and H3K36me3-containing nucleosomes inhibit PRC2 using two distinct mechanisms due to the differences in their positions along the histone H3 tail. These mechanisms of inhibition target two important requirements for the efficient trimethylation of histone H3K27 and support a model in which chromatin regions rich in H3K4me3 or H3K36me3 can act as boundaries to restrict PRC2 function to confined locations in the genome. We also further define important functions for the accessory protein JARID2 in alleviating inhibition by these chromatin modifications that may be critical when large sections of the genome require silencing during embryonic development.

## Results

### PRC2 engagement with the histone H3 tail is reduced by H3K36me3

Histone H3K36me3 decorates gene bodies and interacts directly with RNA polymerase II to regulate transcription elongation^32,38,39^. The deposition of H3K27me3 is prevented at actively transcribed genes through uncharacterized mechanisms, and interestingly, histone H3K36me3 directly inhibits the activity of PRC2 in biochemical assays^5,18,25,26^. Previous cryo-EM structures of PRC2 bound to unmodified nucleosome substrates described the extensive network of interactions that facilitate the placement of K27 at the EZH2 active site for catalysis. The unmodified histone H3K36 was hypothesized to be important for PRC2 activity due to its prominent location at the entry site of the H3 tail binding channel, where it forms electrostatic interactions with the nucleosomal DNA and polar contacts with the CXC domain^5,18^. On the other hand, the PRC2 core has been shown to efficiently monomethylate H3K27 (H3K27me1) in H3K36me3 nucleosomes and, in the presence of JARID2, exhibits some level of higher order methylation on such substrates^5,18,25,26^.

To investigate the molecular basis for the inhibition of PRC2 by chromatin marked with H3K36me3, we sought to determine structures of PRC2 bound to modified nucleosome substrates utilizing a PRC2 complex containing AEBP2 and a construct of JARID2 that included the allosterically activating K116me3 segment **(Figure 1A).** This PRC2 complex (referred to as PRC2_AJ1-450_) was used in previous cryo-EM studies of PRC2 bound to nucleosome substrates and shown to exhibit significantly reduced activity on H3K36me3-modified nucleosomes with respect to unmodified nucleosome substrates, but higher activity than PRC2 complexes lacking JARID2_K116me3 5_. All structural efforts described in this paper utilized streptavidin affinity grids^40,41^, which provide a robust sample preparation strategy to protect the PRC2-nucleosome complexes from the adverse effect of the air-water interface and to enrich for nucleosome-bound PRC2^5^.

**Figure 1.**
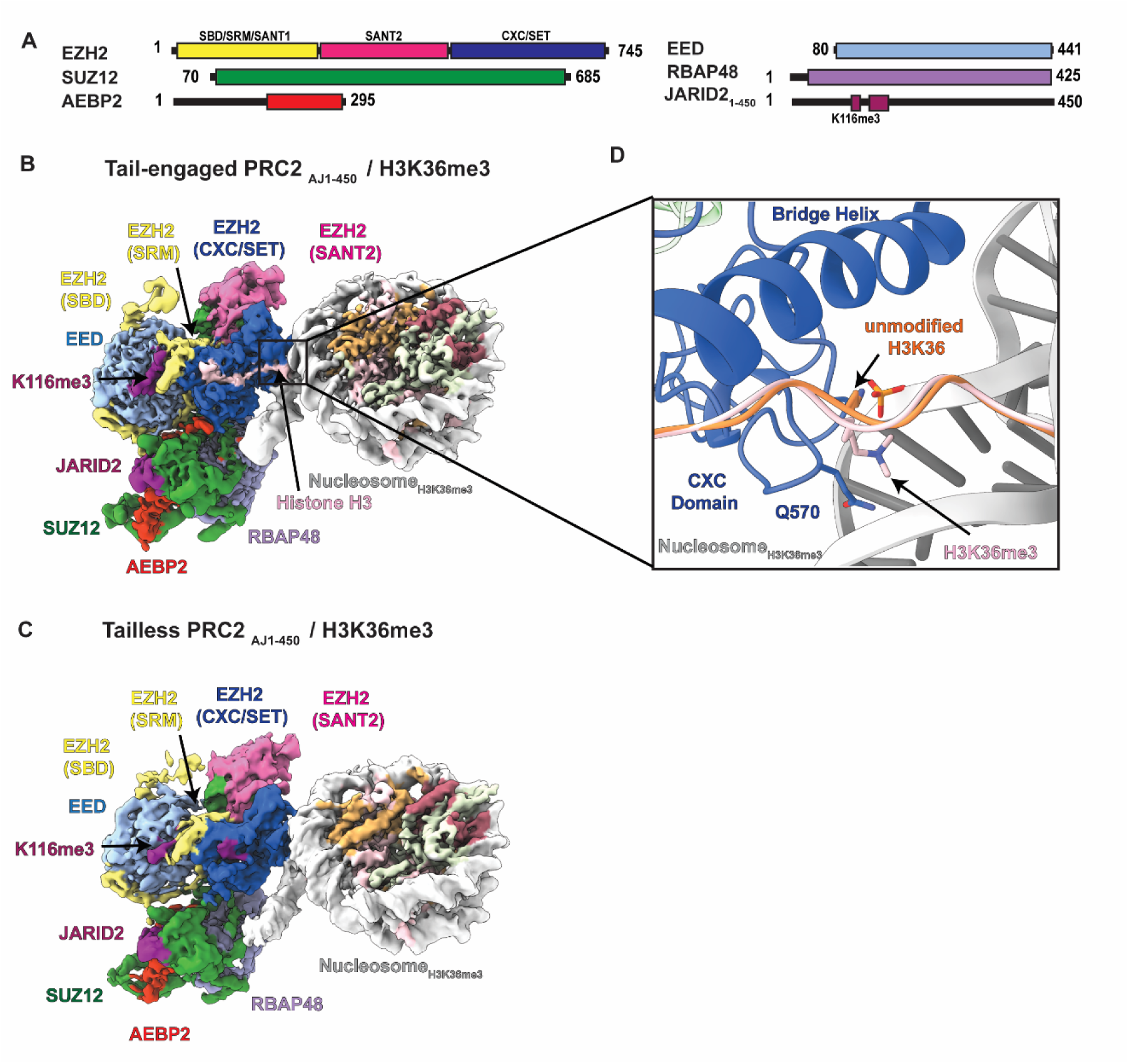
Cryo-EM structures of PRC2_AJ1-450_ bound to H3K36me3 modified nucleosomes. **A)** Schematic representation of protein domains in the PRC2-AEBP2-JARID2 complex used in this work, containing either JARID2_1-450_ or JARID2_119_. **B)** Cryo-EM structure of PRC2_AJ1-450_ bound to a H3K36me3-modified nucleosome in which the H3 tail is engaged by EZH2 and PRC2 is in an allosterically stimulated state. **C)** Cryo-EM structure of PRC2_AJ1-450_ bound to H3K36me3-modified nucleosomes in which the H3 tail is not engaged by EZH2. The structures shown in B) and C) co-exist in the sample. **D)** Comparison of the position of K36 with respect to the nucleosomal DNA in structures of PRC2 bound to unmodified H3K36 (shown in orange; PDB 6WKR) and H3K36me3- modified nucleosomes (shown in pink).

Analysis of the cryo-EM data of PRC2_AJ1-450_ bound to the H3K36me3-modified nucleosome resolved two distinct states at 3.1 Å and 3.5 Å resolution, respectively. In the first state the H3 tail can be seen binding to EZH2 and entering the EZH2 active site (referred to as “tail-engaged”) (**Figure 1B, Figure S1, Figure S2A)**, and in the second state density for the H3 tail is absent (referred to as “tailless”) (**Figure 1C, Figure S1, Figure S2B)**. In both the “tail-engaged” and the “tailless” states, we observe JARID2_K116me3_ bound at the allosteric site on EED. Consequently, as previously observed for activated forms of the complex, the SRM is folded and interacting with the SET-I helix of the active site, and the SBD is in a bent conformation (another conformational feature associated with an active state^6,7,11^) **(Figure 1B-C)**. The presence of H3K36me3, therefore, does not disrupt the allosteric communication between the EED and EZH2 core subunits.

In the “tailless” state, no density is seen extending from the histone core and interacting with the structural elements that channel it to the active side (**Figure 1C)**. There is residual density at the active site that we attribute to a second JARID2 molecule, as previously reported in structural work on JARID2-containing PRC2 complexes lacking histone substrates^6^. The “tail-engaged” state, on the other hand, closely resembles that of our previously published structure of PRC2 bound to a nucleosome containing an unmodified H3K36^5^, with continuous density for the H3 tail along the substrate-binding cavity, and with H3K27 positioned for methylation at the active site, despite the presence of the H3K36me3 modification. We found that in such a state, the H3K36me3 is accommodated by a displacement of the lysine side chain compared to our previous PRC2/nucleosome structure (PDB: 6WKR)^5^ containing unmodified H3K36 (**Figure 1D)**. This small change preserves the majority of contacts between PRC2, the nucleosomal DNA, and the histone H3 tail. The mixture of “tail-engaged” and “tailless” states observed for PRC2 when in the presence of the histone H3K36me3 modification suggests that this mark impacts the efficiency of PRC2 to engage the histone H3 tail. On the other hand, when the histone H3 tail is bound, as captured in the “tail-engaged” state, the H3K36me3 modification does not affect the binding of JARID2 to the EED regulatory site and the allosteric communication between the EED and EZH2 core subunits are not disrupted.

### H3K36me3 modifies the interaction between PRC2 and chromatin

Further comparison of the “tail-engaged” and “tailless” PRC2_AJ1-450_/H3K36me3 states revealed that the “tailless” complex is rotated with respect to the nucleosome interface by approximately 12 degrees **(Figure 2A)**. This rotation results in a different DNA-binding register for both the EZH2 bridge helix **(Figure 2B)** and CXC domain **(Figure 2C)** that offsets the bridge helix by approximately two helical turns (i.e., seven residues). This seven-residue offset results in additional contacts between the bridge helix and nucleosomal DNA, involving residues that otherwise interact directly with the histone H3 tail in the “tail-engaged” state **(Figure 2B)**. Of notice, in the “tail-engaged” state, R504 interacts with the backbone of K36 in the histone H3 tail and R497 forms an electrostatic interaction with the backbone of the nucleosomal DNA. In the “tailless” state, R504 satisfies the DNA contact made by R497 in the “tail-engaged” state, while R497 now makes an additional DNA contact upstream. Similarly, this new binding geometry in the “tailless” state relocates bridge helix residue Q507 to a position where it can no longer interact with the H3 tail as is observed in the “tail-engaged state.” In addition, the CXC domain in the “tailless” state now makes contact with the major groove of the nucleosome DNA at the SHL 6.5 position (while it contacts SHL 7 in the “tail-engaged” state) (**Figure 2C)**. Overall, despite this offset, both the residues involved, and the nature of their interactions are maintained, consistent with the nonspecific nature of the CXC- and bridge helix-mediated DNA interactions.

**Figure 2.**
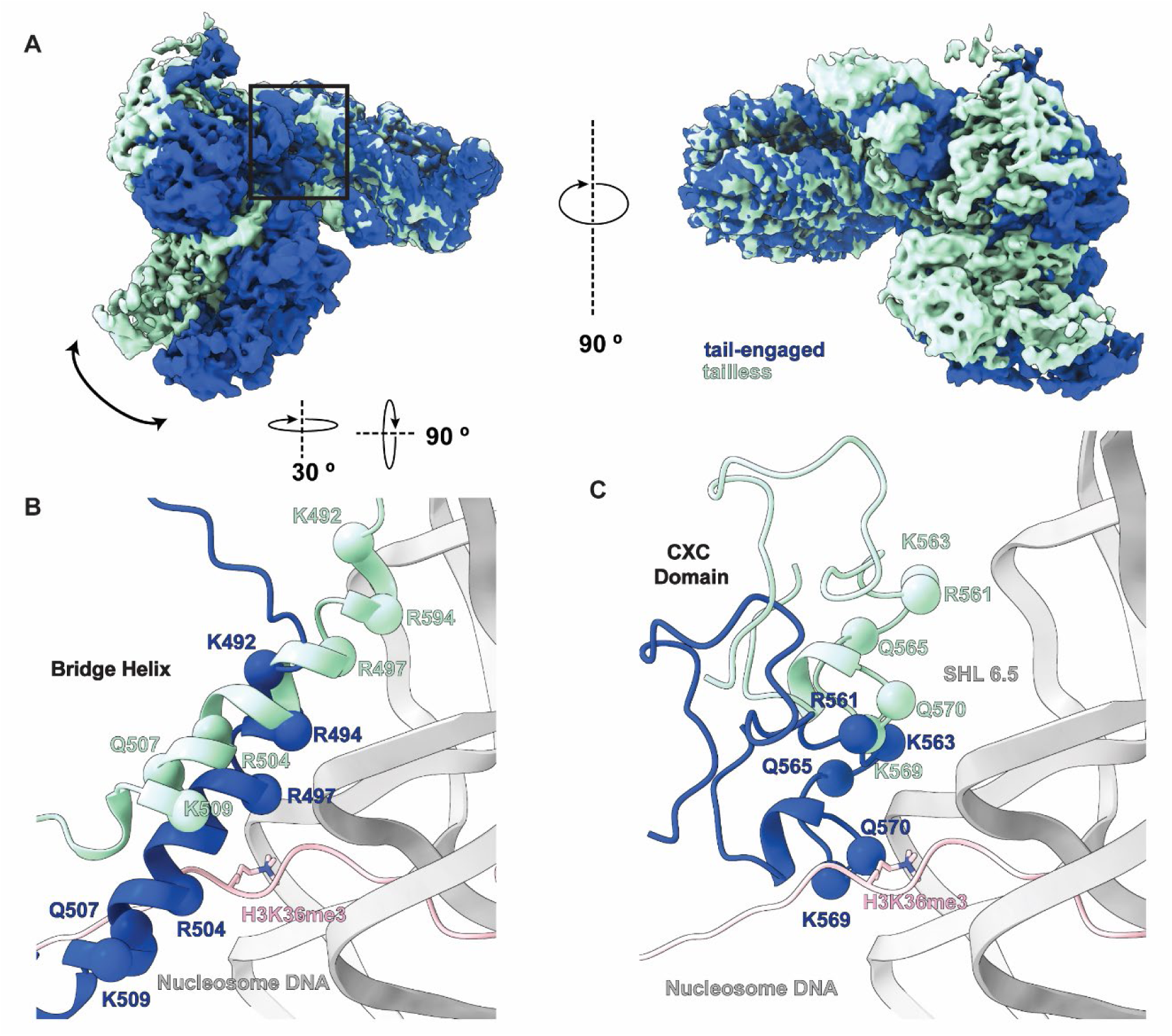
Comparison of tail-engaged and tailless PRC2_AJ1-450_ complexes bound to H3K36me3-modified nucleosomes. **A)** Overlay of the cryo-EM density maps for the co-existing tail-engaged (blue) and tailless (green) PRC2_AJ1-450_ / H3K36me3 structures identified by our analysis. Maps are aligned using the nucleosome to show the relative rotation of PRC2 on the nucleosome surface. **B)** Close up view of the EZH2 bridge helix showing its relative position with respect to the H3 tail (pink) and nucleosomal DNA (grey) for the tail-engaged (blue) and tailless (green) structures. **C)** Close up view of the EZH2 CXC domain showing its relative position with respect to the H3 tail and nucleosomal DNA for the tail-engaged (blue) and tailless (green) structures (in B and C the tail, as seen in the tail-engaged complex, is shown in pink).

In previous structures of PRC2 engaged with nucleosome substrates, the carbonyl group of PRC2 Q570 interacts with the epsilon-amino group of K36 in the histone tail^5,18^. The loss of hydrogen-bonding potential with the CXC domain when H3K36 is trimethylated most likely contributes to the alternative binding geometry that we observe in our cryo-EM data. Importantly, this alternative binding geometry in the presence of H3K36me3 reveals a novel interaction mode of PRC2 with chromatin that is incompatible with catalytic activity and highlights the sensitivity of the H3-tail binding pocket within EZH2 to the surrounding chromatin environment.

### PRC2 interaction with chromatin is variable in the absence of the histone H3 tail

To investigate how PRC2 engages with nucleosomes modified with H3K36me3 in the absence of allosteric activation, we omitted JARID2_K116me3_ from the sample preparation. A minimal construct of JARID2 that contains residues 119-450 has been shown to stabilize the PRC2 core, reduce dimerization of PRC2, and was essential for reaching high resolution in our previous cryo-EM studies of PRC2 complexes^6,17^. Therefore, we included this JARID2 construct in all structural efforts lacking JARID2_K116me3_ described in this paper. Under such conditions, PRC2 has been shown to exhibit further reduced trimethylation activity on nucleosome substrates harboring H3K36me3^5,25^.

Despite much effort, it was not possible to produce a structure of PRC2 lacking JARID2_K116me3_ bound to the H3K36me3 nucleosome due to the flexibility/heterogeneity of the engagement between them. Extensive 2D classification of a large cryo-EM data set showed only fuzzy density bound to nucleosomes **(Figure S3A & Figure S4A)**, suggesting that the positioning of PRC2 with respect to the nucleosome is variable (i.e., there is no fixed register of PRC2 on the nucleosome) under these conditions. As a control, we could obtain reconstructions of the PRC2 complex lacking JARID2_K116me3_ when it was bound to unmodified nucleosomes from just a few hundred micrographs (**Figure S3B**). These results strongly suggest that the increased mobility observed at the PRC2/nucleosome interface is caused by the presence of the H3K36me3 modification and its negative effect on the engagement of PRC2 with substrate H3 tails.

To assess how the histone H3 tail impacts the interaction of PRC2 with chromatin, we performed electromobility shift assays of PRC2 complexes including or lacking JARID2_K116me3_ in the presence of unmodified nucleosomes, H3K36me3-containing nucleosomes, or H3Δ38-containing nucleosomes that lack the H3 residues that interact with EZH2. We found that PRC2 complexes bound to all three nucleosome substrates with similar affinity and independently of the presence of JARID2_K116me3_ **(Figure S4A-D)**. Furthermore, cryo-EM analysis of PRC2_AJ119-450_ bound to nucleosomes lacking the histone H3 tail (N-terminally truncated at residue 38, H3Δ38) closely resembled that obtained for the H3K36me3-modified nucleosome, further supporting that in the absence of H3 tail engagement with EZH2, the PRC2/nucleosome interaction is highly dynamic/variable and difficult to visualize by cryo-EM **(Figure S3C and Figure S4B)**. These results suggest that although the histone H3 tail contributes minimally to the affinity of PRC2 for nucleosomes, which is dominated by electrostatic interactions with nucleosomal DNA^23,42^, the histone H3 tail is important for the functional engagement of PRC2 with chromatin. Several studies suggest that targeting of PRC2 across the genome may be decoupled from its methyltransferase activity^12,43,44^. Our “tailless” PRC2-H3K36me3 state now shows a variable interaction of PRC2 with chromatin that is not productive for activity. This variability highlights a way in which PRC2 can engage chromatin without performing its catalytic function.

In summary, our results are consistent with a mechanism in which the histone H3K36me3 modification reduces productive engagement of the histone H3-tail with PRC2 and promotes dynamic interactions with chromatin that are incompatible with the trimethylation of histone H3K27 by PRC2 at actively transcribed gene bodies rich in this modification. This effect is partially overcome in the presence of JARID2_K116me3_, which interacts with both EED and EZH2 and thus stabilizes the catalytic lobe of the complex in a way that helps it retain the histone H3 tail in the active site.

### H3K4me3 binds to the EED allosteric site

Histone H3K4me3 localizes to actively transcribed promoters, where it has been demonstrated to interact directly with the transcription initiation machinery and recently shown to regulate promoter-proximal pausing^45–48^. In vitro biochemical assays show that the H3K4me3 modification directly inhibits the activity of PRC2, but only when it is present in cis on nucleosome substrates^25^. To investigate how H3K4me3 inhibits PRC2 function, we obtained a 3 Å resolution structure of PRC2 in the absence of JARID2_K116me3_ and bound to an H3K4me3-modified nucleosome **(Figure S5 & Figure S6A)**. Overall, this complex resembles other PRC2/nucleosome structures, in which the H3 tail is clearly visible entering the active site. Unlike the interactions with H3K36me3-modified nucleosomes, we were unable to identify a population of particles lacking tail engagement. Not surprisingly, given the absence of JARID2_K116me3_ as an allosteric activator, there is no density for the SRM, and the SBD is in an extended conformation, both indicative of an unstimulated state (**Figure 3A)**. However, we do observe a methylated lysine bound to the aromatic cage of EED. We propose that this density corresponds to the H3K4me3 modification (**Figure 3B**), the source of methylated peptide in the sample. Indeed, the resolution of the PRC2_AJ119-450_/H3K4me3 structure allowed us to confidently build a peptide into the unassigned density with a sequence matching that of the N-terminus of the H3 tail **(Figure S7A-B)** that agrees with a previous crystal structure of EED bound to a H3K4me3-containing peptide^49^. Although the affinity of EED for the H3K4me3 peptide was previously reported to be low, it is important to notice that ours is the first structure reporting the interaction of H3K4me3 with EED in the context of the nucleosome, the relevant PRC2 substrate. To further eliminate the possibility that this peptide originated from the sample purification, we also determined the structure of PRC2_AJ119-450_ bound to an unmodified nucleosome substrate. In this structure **(Figure S7C & Figure S8)**, and as in a previous reconstruction of PRC2/nucleosome also lacking JARID2K119me3 **(Figure S7D)**^17^, the EED aromatic cage is vacant. We therefore assigned the extra density in the PRC2_AJ119-450_/H3K4me3 structure to the H3 peptide encompassing K4me3.

**Figure 3.**
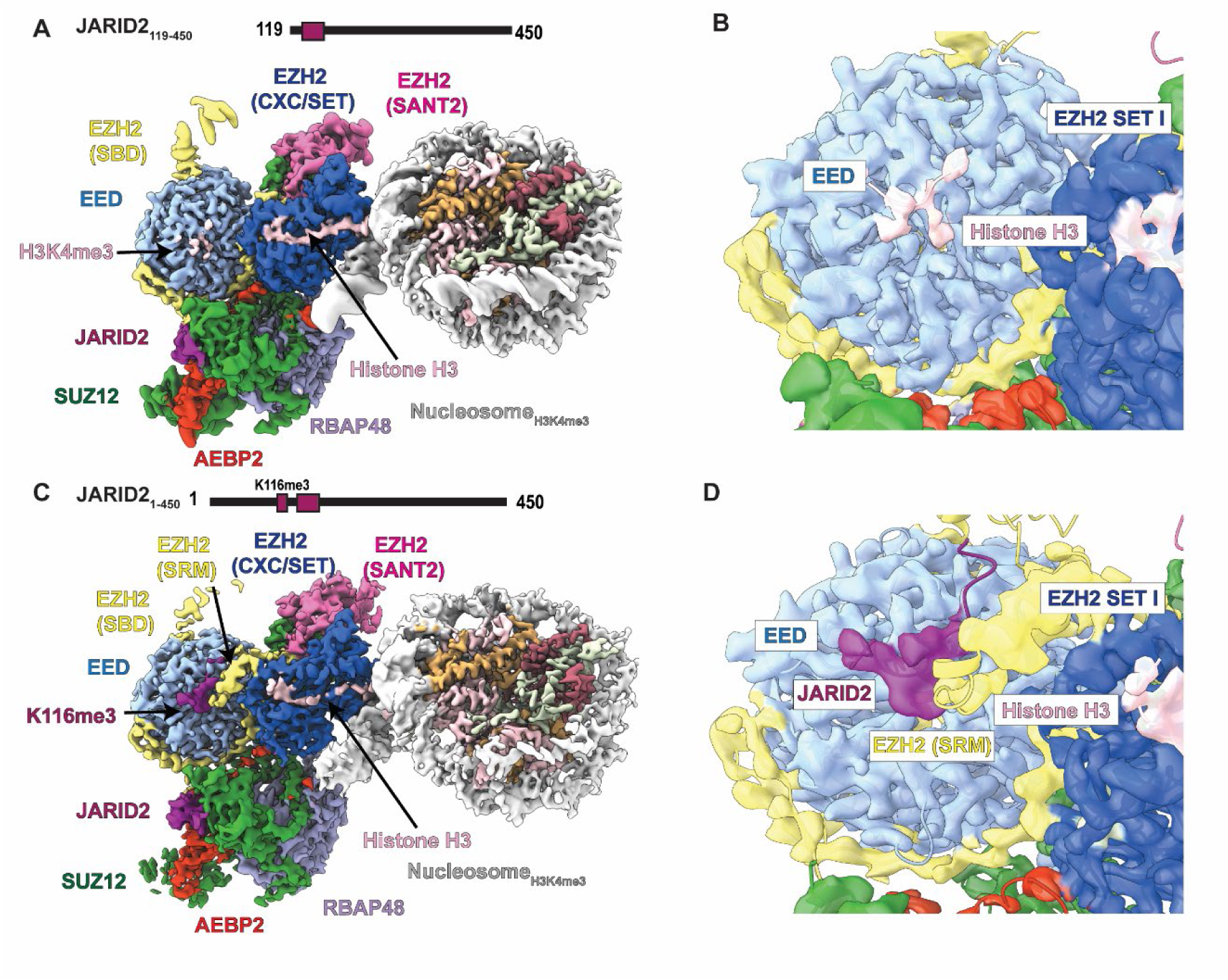
Cryo-EM structures of PRC2 bound to H3K4me3-modified nucleosomes. **A)** Cryo-EM structure of PRC2_AJ119-450_ bound to H3K4me3-modified nucleosomes. **B)** Close up view of the cryo-EM density showing the N-terminus of histone H3 containing H3K4me3 bound to EED. Map is shown at threshold 0.487. For lower threshold refer to Figure S7A. **C)** Cryo-EM structure of PRC2 _AJ1-450_ bound to H3K4me3-modified nucleosomes. **D)** Close up view of cryo-EM density showing JARID2_K116me3_ bound to EED and the folding of the SRM helix. Map is shown at threshold 0.25.

### JARID2 can compete with H3K4me3 for the EED allosteric site

Previous biochemical data showed that PRC2 can be activated on H3K4me3-modified nucleosome substrates in the presence of high local concentrations of allosteric activator (either excess H3K27me3 peptide or PRC2 complexes copurified with methylated JARID2)^5,25^. We therefore determined the structure of PRC2_AJ1-450_, thus including JARID2_K116me3_, bound to an H3K4me3- modified nucleosome at 3.5 Å resolution **(Figure S9)**. In this structure, we observed an activated state of PRC2, with density consistent with JARID2_K116me3_ occupying the EED regulatory site, bent SBD, and folded SRM **(Figure 3C-D & Figure S10A).** Importantly, we could not locate the histone H3K4me3 modification elsewhere in this reconstruction, supporting that JARID2_K116me3_ and H3K4me3 compete for the same binding site.

Although the activated state was prevalent in our data, careful sorting for occupancy of the SRM led us to identify a second, more rare state (resolution limited to 8 Å), in which the SRM is unfolded, and the SBD is extended, although the EED regulatory site remains occupied **(Figure S10B)**. These features correspond to a non-activated state of PRC2, similar to that obtained for the PRC2_AJ119-450_/H3K4me3 structure. The existence of both allosterically activated and inactivated PRC2 states in the cryo-EM data, agrees with previous biochemical observations concerning activity under conditions including allosteric activators, and suggests that JARID2 and H3K4me3 are mutually exclusive binders of the EED regulatory site.

### H3K4me3 acts as an allosteric antagonist

The recognition of methylated-lysines by EED occurs through a classical hydrophobic cage. Previous studies indicate that sequence variation among EED-binding peptides is tolerated by subtle alternative binding modes involving residues within EED and EZH2^7,9,49^. Our study shows that histone H3K4me3 interacts with the aromatic cage in a similar manner to other EED-binding peptides and is further stabilized by interactions between histone residue H3T6 and the hydrophobic pocket defined by I363, Q382, and A412 of EED **(Figure 4A)**. To reconcile our finding that H3K4me3 binds to the EED regulatory site but fails to activate PRC2, we further compared its potential to interact with the EZH2 SRM with that of known allosteric activators of PRC2.

**Figure 4.**
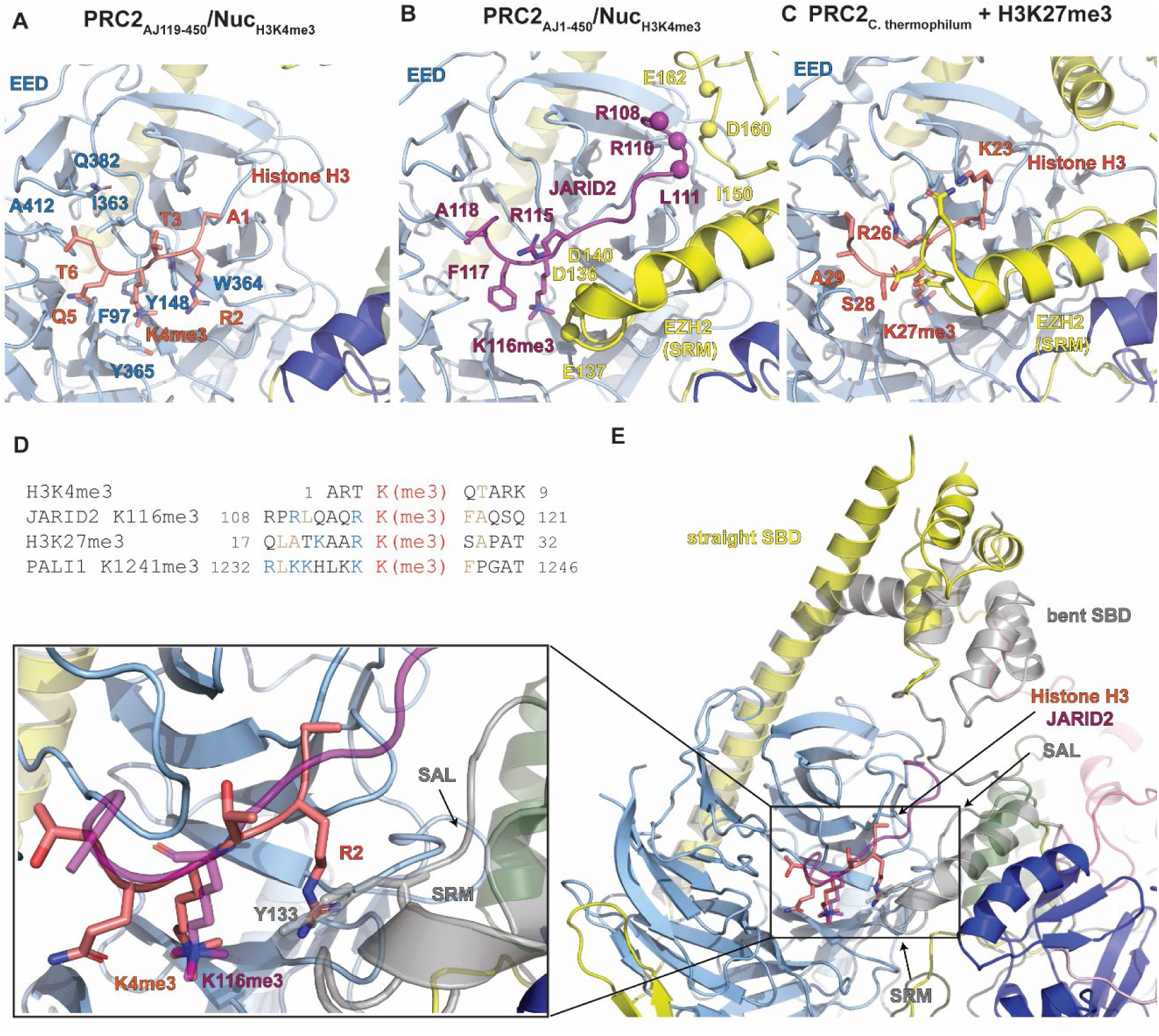
H3K4me3 acts as an allosteric antagonist by binding to EED. **A)** Interactions between the region of histone H3 (pink) around the H3K4me3 modification and the aromatic cage of EED (light blue) **B)** Interactions between JARID2 (magenta), EED (light blue), and EZH2 (yellow for the SRM and SAL, and darker blue for the SET domain) in an allosterically activated PRC2. The coordinates used were from the PRC2_AJ1-450_/H3K4me3 structure obtained in this study and shown in Fig. 3C. JARID2 R115 interacts with E137 and D140 of the EZH2 SRM. JARID2 R108 and R110 are positioned to interact with E162 and D160 of EZH2, while JARID2 L111 and EZH2 I150 are involved in hydrophobic contacts. **C)** Interactions between the peptide around H3K27me3 (pink) and EED (light blue) and EZH2 (yellow for the SRM and SAL, and darker blue for the SET domain). in the coordinates used are those for the X-ray crystal structure of an activated PRC2 catalytic lobe from *C. thermophilum* (PDB: 5KJH)^7^. Histone H3R26 (similarly to JARID2 R115) interacts with negatively charged residues in the EZH2 SRM, and histone H3R23 establishes additional contacts with the SRM. **D)** Sequence alignment with respect to the PRC2- methylated lysine for: the N-terminal region of the histone H3 tail around K4, JARID2 around K116, H3 around K27, and PALI1 around K1241. The PRC2-modified lysine is colored in red, residues that are involved in hydrophobic contacts are colored in tan, and residues involved in electrostatic interactions are colored in blue. **E)** Overlay of structures shown in panels **A** (colored) and **B** (grey/transparent) showing that histone H3R2 of H3K4me3 clashes with the SAL.

Our structures containing JARID2 K116me3 show that JARID2 is positioned to form several hydrophobic and electrostatic interactions with residues of the EZH2 SRM (**Figure 4B**), as also described in previous cryo-EM and crystallographic studies of JARID2-containing PRC2 complexes^5,6,11^. Similar electrostatic interactions were also observed in a crystal structure of PRC2 bound to a H3K27me3-containing peptide (**Figure 4C),** and biochemical perturbation experiments support the importance of these SRM residues for the stimulation of PRC2 activity^7,12^. Notably, the interactions with the EZH2 SRM created by JARID2 or H3K27me3 involve residues that are N-terminal to the methylated lysine. Other activators, such as PALI1 K1241me3, contain sequence similarities at equivalent positions (specifically the positively charged residue in the −1 position), while these residues are not conserved or cannot exist in H3K4me3 due to its location proximal to the N-terminus of the histone H3 protein **(Figure 4D)**. Additionally, histone H3R2, the only positively charged residue that is N-terminal to histone H3K4me3, sterically clashes with the SANT activation loop (SAL) in EZH2 **(Figure 4E)** that is required for stimulation through EED^7^ and, as a consequence, density for the SAL is absent in our PRC2_AJ119_/H3K4me3 structure.

Overall, the absence of contacts observed between residues around histone H3K4me3 and EZH2 may explain why, in spite of its ability to engage with the aromatic cage of EED, it fails to stabilize the SRM for stimulation of EZH2 activity seen for other EED-binders that have been characterized as allosteric activators. An alternative (or additional) explanation that we are unable to exclude, is that residues 7 through 23 of the histone H3 tail, which are invisible in our structure of PRC2_AJ119-450_/H3K4me3 **(Figure 3C)**, could themselves sterically block the folding of the SRM when H3K4me3 is engaged by the allosteric site. In conclusion, our structural analysis leads us to propose that the abundance of H3K4me3 localized on actively transcribed chromatin acts as allosteric antagonist to decrease PRC2 activity at these genomic locations.

## Discussion

The combination of both positive and negative PRC2 regulatory mechanisms is required to fine tune PRC2 activity to restrict the H3K27me3 repressive mark to specific genomic locations. This fine tuning is achieved via the activation and inhibition of the complex through different mechanisms involving crosstalk between PRC2, its accessory subunits, and existing marks on chromatin **(Figure 5)**. Both H3K4me3 and H3K36me3 are thought to serve as physical barriers for the spreading of H3K27me3. In support of this model, the depletion of either H3K4me3 or H3K36me3 methyltransferase machinery results in the redistribution of PRC2 across the genome and the invasion of H3K27me3 into domains that are decorated with H3K4me3 or H3K36me3 under normal conditions ^36,50,51^.

**Figure 5.**
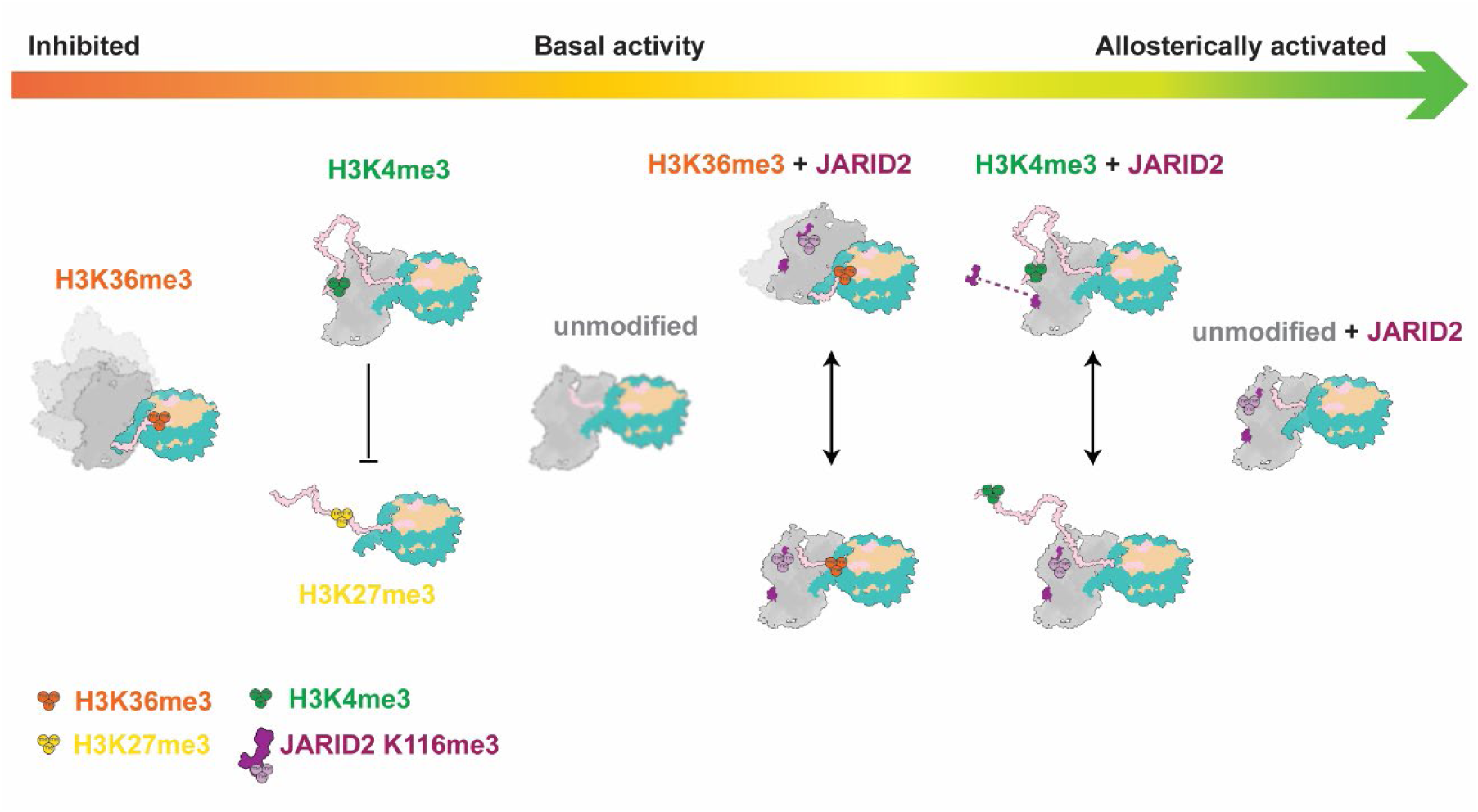
A structural model for tuning the catalytic activity of PRC2 in vitro. Trimethylation of histone H3K27 by PRC2 can be inhibited by H3K36me3 and H3K4me3 histone modifications and activated by H3K27me3 or JARID2 K116me3. The interplay between these regulators is a gradient of PRC2 activity (red to green arrow). From left to right: in the absence of JARID2, H3K36me3 prevents efficient tail engagement and results in the loss of a well-defined register between PRC2 and the nucleosome substrate, as depicted as the blurry model of PRC2.; histone H3K4me3 engages the allosteric site and competes with substrates that are already modified with H3K27me3; PRC2 on unmodified substrates in the absence of JARID2 has a basal level of activity, while its EED allosteric site remains open for possible engagement with nearby nucleosomes with H3K27me3; in the presence of JARID2, PRC2 is allosterically stimulated, resulting in increased H3K27me3 activity (right most); histone H3K36me3 and H3K4me3 reduce PRC2 activity through the same mechanisms described above, however, methylated JARID2 stabilizes the catalytic lobe of PRC2 and facilitates some tail-engagement in the presence of H3K36me3, and it can compete with H3K4me3 for the EED allosteric site.

In this study, we investigated the molecular mechanisms by which the histone PTMs H3K36me3 and H3K4me3, which are present in actively transcribed genes, directly inhibit the activity of PRC2. We find that H3K36me3 and H3K4me3 act through two distinct mechanisms due to their unique and distinct locations on the N-terminal histone H3 tail, and target two important requirements for the higher order methylation of histone H3K27, respectively: 1) the arrangement between PRC2, chromatin, and the histone H3 tail that is conducive to efficient substrate engagement, and 2) the activation of the catalytic SET domain through the EED-EZH2 regulatory axis. These two features permit the histone H3 tail to be retained in the active site for higher order methylation states (di- and tri-methylation), which are the rate limiting steps during PRC2 catalysis ^52^. Both mechanisms of inhibition identified in this study provide potential avenues to break the auto-activating positive feedback loop established through EED-EZH2 and prevent the spreading of the H3K27me3 silencing mark into actively transcribed chromatin.

Histone H3K36 is adjacent to the nucleosome core particle, positioned at the entry site of the histone H3 tail in its path to the PRC2 active site. PRC2 interaction with nucleosomes involve two DNA-binding surfaces on EZH2, the bridge helix and the CXC domain, which directly interact with the unmodified H3K36 sidechain^4,5,18^. Our structural data show that the trimethylation of H3K36 interferes with these interactions, resulting in the loss of a well-defined register for PRC2 nucleosome interactions that is required for effective H3 tail engagement and methylation. We propose that the proper, stable engagement of the histone H3 tail, although not the driver for the interaction between PRC2 and chromatin, is necessary to stabilize PRC2 on chromatin in a way that poises it for activity.

Histone H3K4, on the other hand, is located close to the N-terminus of the flexible histone H3 tail, twenty-three residues upstream from the residue targeted by PRC2 (H3K27) and usually invisible in cryo-EM structures of PRC2 bound to nucleosome substrates. In our structure of PRC2 bound to H3K4me3-modified nucleosomes, we see that H3K4me3 can engage with the EED aromatic cage and occupy the allosteric site of EED. The EED-EZH2 allosteric activation mechanism has been studied extensively and is central to PRC2 function and its spreading of the H3K27me3 repressive mark^9^. The known EED-binders are methylated peptides generated by the activity of EZH2, and include the accessory proteins JARID2 (in PRC2.2) and PALI1 (in PRC2.1), the histone H3K27me3 itself, and, more recently identified, the EZH2 automethylation loop^9,13,17,53^. Because these methylated peptides take advantage of the same regulatory site, they are mutually exclusive and therefore must be utilized in specific cellular contexts. Unlike H3K27me3, JARID2 K116me3, or PALI1, which allosterically stimulate PRC2, we show here for the first time that the engagement of H3K4me3 with EED is unique in that it fails to stabilize the SRM helix and stimulate PRC2 activity. At actively transcribed promoters, in which chromatin is highly decorated with H3K4me3 and low in H3K27me3, the high local concentration of this mark may out-compete activators for the EED regulatory site, thus acting as an allosteric antagonist.

PRC2 is the sole writer in mammals of mono-, di-, and trimethylation of histone H3K27, with only the latter being associated with gene silencing^44^. Biochemical experiments have shown that both H3K4me3 and H3K36me3 have minimal effect on the monomethylation of H3K27^18,25^, and indeed, H3K27me1 is broadly found in the genome, including at actively transcribed regions,^44,50,54^. Higher order methylation, however, is prevented in those regions and appears to require both activation through the EED-EZH2 axis and stable engagement with the dynamic histone H3 tail. Dynamic expression levels of accessory PRC2 subunits such as JARID2 throughout development and across tissues may play a role in alleviating inhibition by H3K4me3 and H3K36me3^2^. For example, the increased expression of JARID2 may favor the engagement of the EED regulatory site with activating methyl-lysine bearing peptides to start spreading the H3K27me3 repressive mark. In support of this model, the JARID2 cofactor is required during differentiation, when large sections of the genome require silencing, yet is dispensable in undifferentiated embryonic stem cells ^13,55,56^. Additionally, our studies now show that the presence of stoichiometric, methylated JARID2 gives rise to stimulated PRC2 complexes engaged with H3K4me3-containing nucleosomes, and are consistent with in vitro biochemical experiments showing that PRC2 activity on H3K4me3-modified substrates can still be enhanced by JARID2 or H3K27me3^5,25^. Similarly, the presence of JARID2_K116me3_, also allowed us to obtain structures of stimulated PRC2 complexes bound to H3K36me3 modified substrates. The presence of methylated JARID2 in the EED results in the global stabilization of the catalytic lobe of PRC2 and enabled us to capture a dynamic state in which the H3K36me3-containing tail is accommodated by the subtle repositioning of the H3K36 sidechain. A number of well-characterized roles in PRC2 regulation have already been assigned to JARID2: (1) residues 138-166 of JARID2 contribute to the stability to the PRC2.2 complex^6^; (2) JARID2_K116me3_ functions as a strong allosteric activator of PRC2 that can perform de novo H3K27me3^13,56^; and (3) its ubiquitin-interaction motif recruits and stabilizes PRC2 on H2AK119Ub- modified chromatin^5,15^. Each of these activity-promoting functions could contribute to rescuing the inhibitory mechanisms imparted by either H3K4me3 or H3K36me3 and thus facilitate the establishment of new heterochromatin domains.

The structures reported here provide mechanistic insight into the tight regulation of PRC2, involving a complex crosstalk between the PRC2 core, its accessory subunits, and the preexisting chromatin environment. Uncovering these and potential new mechanisms of regulation is essential for an understanding of how the polycomb group proteins promote the correct gene silencing to safeguard developmental processes and to maintain cell identity.

## Materials and Methods

### Expression and purification of PRC2.2 complexes

PRC2 complexes containing AEBP2 and either JARID2_1-450_ or JARID2_119-450_ were cloned, expressed, and purified as previously described^6^. Briefly, EZH2, EED, RBAP48, SUZ12 and strep-tagged-GFP fusions of AEBP2 and JARID2 were cloned into the Macrobac system for baculovirus expression in Sf9 insect cells. Cells were resuspended in lysis buffer (50 mM HEPES pH 7.9, 250 mM NaCl, 0.5 mM TCEP, 10% glycerol, 0.1% NP40) and supplemented with a EDTA-free protease inhibitor cocktail, leupeptin, pepstatin A, aprotinin, and benzonase. Cells were lysed by sonication and cleared by centrifugation at 14,500 rpm for 45 minutes. Supernatant was incubated with Strep-Tactin Superflow Plus resin overnight, washed with buffer containing 1 M NaCl, and eluted with 10 mM desthiobiotin. The eluate was subjected to TEV protease cleavage followed by size exclusion chromatography with a Superose 6 Increase 3.2/300 equilibrated with 50 mM HEPES pH 7.9, 150 mM KCl, 0.5 mM TCEP, 10% glycerol. The purified complex was flash frozen in liquid nitrogen and stored at −80 °C as single-use aliquots.

### Nucleosome purification

For use in both cryo-EM and EMSAs experiments, human nucleosomes containing unmodified H3, H3K4me3, or H3K36me3 were purchased from Epicypher with biotinylated DNA containing the following sequence:

GGACCCTATACGCGGCCGCCCTGGAGAATCCCGGTCTGCAGGCCGCTCAATTGGTCGTAGACAGCTCTAGCACCGCTTAAACGCACGTACGCGCTGTCCCCCGCGTTTTAACCGCCAAGGGGATTACTCCCTAGTCTCCAGGCACGTGTCAGATATATACATCCTGTGCCGGTCGCGAACAGC GACC3’

Human octamers lacking the H3 tail were purchased from The Histone Source and reconstituted into nucleosomes by standard protocols. Biotinylated DNA containing the following sequence

ATATCTCGGGCTTATGTGATGGACCCTATACGCGGCCGCCCTGGAGAATCCCGGTGCCGAGGCCGCTCAATTGGTCGTAGACAGCTCTAGCACCGCTTAAACGCACGTACGCGCTGTCCCCCGCGTTTTAACCGCCAAGGGGATTACTCCCTAGTCTCCAGGCACGTGTCAGATATATACATCCTGTGCATGTATTGAACAGCGACTCGGGATAT

was amplified by PCR and purified over a monoQ column and ethanol precipitation before being resuspended in high salt buffer (10 mM Tris-HCl, pH 7.5, 2 M KCl, 1 mM EDTA, 1 mM DTT). 4.4 μM octamer was mixed with 4 μM biotinylated DNA and subject to slow dialysis over 16 hours into low salt buffer (10 mM Tris-HCl, pH 7.5, 250 mM KCl, 1 mM EDTA, 1 mM DTT) before being dialyzed into nucleosome storage buffer (20 mM Tris-HCl, pH 7.5, 1 mM EDTA, 1 mM DTT).

### Cryo-EM grid preparation

All cryo-EM samples were prepared using streptavidin affinity grids that were fabricated in-house as previously described^40,41^. PRC2/nucleosome complexes were assembled by incubating 100 nM nucleosome with 500 nM PRC2 in cryo buffer (50 mM HEPES pH 7.5, 50 mM KCl, 0.5 mM TCEP, 100 μM SAH). Sample was applied to rehydrated streptavidin affinity grids and incubated for 3-5 minutes at room temperature in a humidity chamber. Following incubation, grids were washed with freezing buffer (50 mM HEPES pH 7.5, 50 mM KCl, 0.5 mM TCEP, 4% trehalose, 0.01% NP40). Excess buffer was manually blotted away and 4 μL of freezing buffer was applied before transferring grids to the Leica GP2 automated plunger. Grids were blotted for 4-5 s using the Leica GP2 blot sensor before plunging into liquid ethane.

### Data collection and processing

High-resolution data sets were collected on a Titan Krios G3i microscope equipped with a Gatan Quantum energy filter (slit width 20 eV) at the University of California, Berkeley QB3 Cal-Cryo facility or at the Stanford-SLAC Cryo-EM Center (S2C2).

Movies from all datasets were motion corrected and dose weighted using MotionCor2^57^ and then the streptavidin lattice was removed from the images using MATLAB^40^. CTF estimation was performed using CTFfind4^58^.

To obtain the reconstruction of PRC2_AJ119-450_ bound to H3K4me3-modified nucleosomes, 22,000 raw movies of 50 frames were collected for dataset 1 and 16,585 raw movies were collected for data set 2. Both data sets were collected using super resolution with a pixel size of 0.525 Å/pix, with a total dose of 50e-/ Å^2^, and defocus ranged between −0.8 and −1.8 μm. ∼6 million particles were picked for each dataset using a trained convolutional neural network in CRYOLO^59^. Initial 2D classification and 3D heterogeneous refinement were performed in CryoSPARC^58^ . Both datasets were merged and imported into RELION^60^ for 3D classification without alignment. CTF refinement was performed to correct for beam tilt, per-particle defocus, and per-micrograph astigmatism. CTF-refined particles were imported into CryoSPARC for local refinement. Maps were filtered by local resolution using manually adjusted B-factors to prevent over-sharpening and the resulting density map was used for modeling.

For PRC2_AJ1-450_ bound to H3K4me3-modified nucleosomes, 15,961 raw movies were collected using super resolution with a pixel size of 0.43 Å/pix. ∼5.6 million particles were picked using template picker in CryoSPARC with 2D classes generated from a smaller subset of particles picked with the CryoSPARC blob picker. Initial 2D classification and 3D Heterogeneous refinement were performed in CryoSPARC. Particles were imported into RELION for 3D classification without alignment. Particles were then subjected to focused classification around the EED/SRM region to identify a subset of particles lacking density for the SRM.

To obtain the reconstruction of PRC2_AJ1-450_ bound to H3K36me3 modified nucleosomes, 14,796 raw movies of 50 frames were collected with super resolution pixel size of 0.525 Å/pix, with a total dose of 50e-/ Å^2^, and defocus range between −0.8 and −1.8 μm. ∼4.3 million particles were picked for each dataset using a trained convolutional neural network in CRYOLO^59^. Initial 2D classification and 3D classification was performed in RELION. Focused classification was performed around the nucleosome and EZH2 SET domain to sort for a state that had density for the histone H3 tail. Focused classification was then performed on the PRC2 top lobe. The tailless class of particles was obtained through heterogeneous refinement in CryoSPARC. CTF refinement was performed for beam tilt, per-particle defocus, and per-micrograph astigmatism for tail engaged and tailless particle stacks. CTF-refined particles were imported into CryoSPARC for non-uniform and local refinements.

For PRC2_AJ119-450_ bound to unmodified nucleosomes, 9,001 raw movies were collected in super resolution with a pixel size of 0.43 Å/pix,. Particles were picked using template picker in CryoSPARC and subjected to several rounds of 2D classification and heterogeneous refinement.

For PRC2_AJ119-450_ bound to H3K36me3 modified nucleosomes, 21,177 raw movies were collected in super resolution with a pixel size of 0.43 Å/pix. Particles were picked using template picker in CryoSPARC and subjected to many rounds of 2D classification to bring out density for PRC2. Only after 11 rounds of 2D classification could PRC2 be observed in the sample. Ab inito reconstruction failed, likely due to lack of views that were sorted out in the extensive cleaning of the data by 2D classification. A second data set of 1,331 movies was collected on a Talos Arctica with super resolution and a pixel size of 0.57 Å/pix, total dose of 50e-/ Å^2^, and defocus range between −0.8 and −1.8 μm. Particles were subjected again to extensive 2D classification to bring out density for PRC2. An ab inito reconstruction was obtained, but PRC2 could not be resolved (data not shown). On the same day, 441 movies were collected for PRC2_AJ119-450_ bound to unmodified nucleosomes. Particles were picked using template picker in CryoSPARC. 2D classification and ab inito reconstruction were performed in CryoSPARC and a low resolution map was obtained containing density for both the unmodified nucleosome and PRC2.

For the sample containing PRC2_AJ119-450_ bound to H3Δ38 nucleosomes, 1,807 raw movies were collected on a Talos Arctica in super resolution with a pixel size of 0.57 Å/pix, total dose of 50e-/ Å^2^, and defocus range between −0.8 and −1.8 μm. Particles were subjected to extensive 2D classification to bring out density for PRC2. An ab inito reconstruction was obtained, but PRC2 could not be resolved.

### Model Building

For all PRC2/nucleosome structures obtained in this study, we used our previously reported structure of PRC2 bound to an ubiquitylated nucleosome (pdb 6WKR) ^5^ as initial model. Coordinates were adjusted using flexible fitting in Isolde v1.5^61^ in UCSF ChimeraX v1.5^62^ and COOT^63^. Models were then iteratively refined and adjusted using PHENIX^64^ and COOT.

### Electrophoretic mobility shift assay (EMSA)

The PRC2/nucleosome reactions were prepared varying the concentration of PRC2 between 0 and 400 nM with 50 nM of either unmodified, H3K4me3, H3K36me3, or H3Δ38 nucleosomes in cryo buffer (50 mM HEPES pH 7.5, 50 mM KCl, 0.5 mM TCEP). Reactions were incubated for five minutes at room temperature. Glycerol was added to the reaction to a final concentration of 5% just before samples were loaded onto a 5% native TBE gel in 0.5X TBE buffer. Gels were stained with SYBR™ Gold.

## Data availability

Cryo-EM maps and fitted models have been deposited in the Electron Microscopy Data Bank (EMDB) and the Protein Data Bank (PDB) under the accession numbers EMD-43361 EMD-43373 EMD-43362 EMD-43363 EMD-43357 EMD-43358 EMD-43359 EMD-43360 and PDB 8VNV 8VOB 8VNZ 8VO0 8VMI 8VMJ 8VML 8VMN. Corresponding accession codes for each structure can be found in Supplementary Table 1.

## Acknowledgments

We acknowledge Dan Toso and Jonathan Remis for microscope support at the Cal-Cryo QB3- Berkeley facility and Paul Tobias, Abhiram Chintangal and Kurt Stine for providing computational support. We thank Alison Killilea at the University of California, Berkeley Cell Culture Facility for providing insect cell cultures. Some of this work was performed at the Stanford-SLAC Cryo-EM Center (S2C2), which is supported by the National Institutes of Health Common Fund Transformative High-Resolution Cryo-Electron Microscopy program (U24 GM129541). The content is solely the responsibility of the authors and does not necessarily represent the official views of the National Institutes of Health. The authors would also like to thank the following S2C2 personnel for their invaluable support and assistance: Patrick Mitchell, Ian Fries, and Lisa Dunn. T.C. was supported by the National Institute of General Medical Sciences molecular biophysics training grant GM-08295 and the National Science Foundation graduate research fellowship program under grant number DGE 2146752. This work was funded through the National Institute of General Medical Sciences grant R35-GM127018 awarded to E.N. E.N. is a Howard Hughes Medical Institute Investigator.

## EXTENDED DATA

**Supplementary Table 1.**
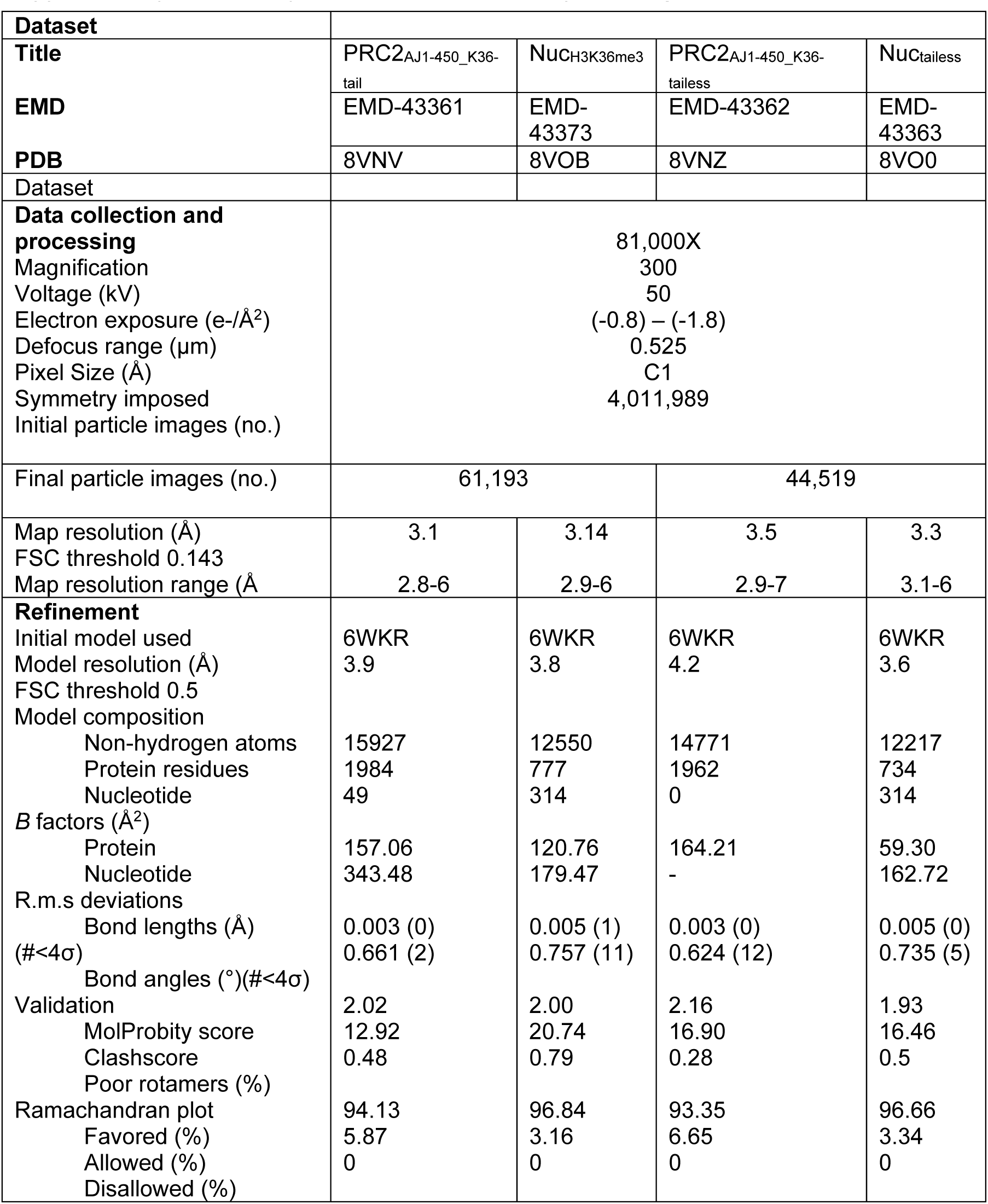
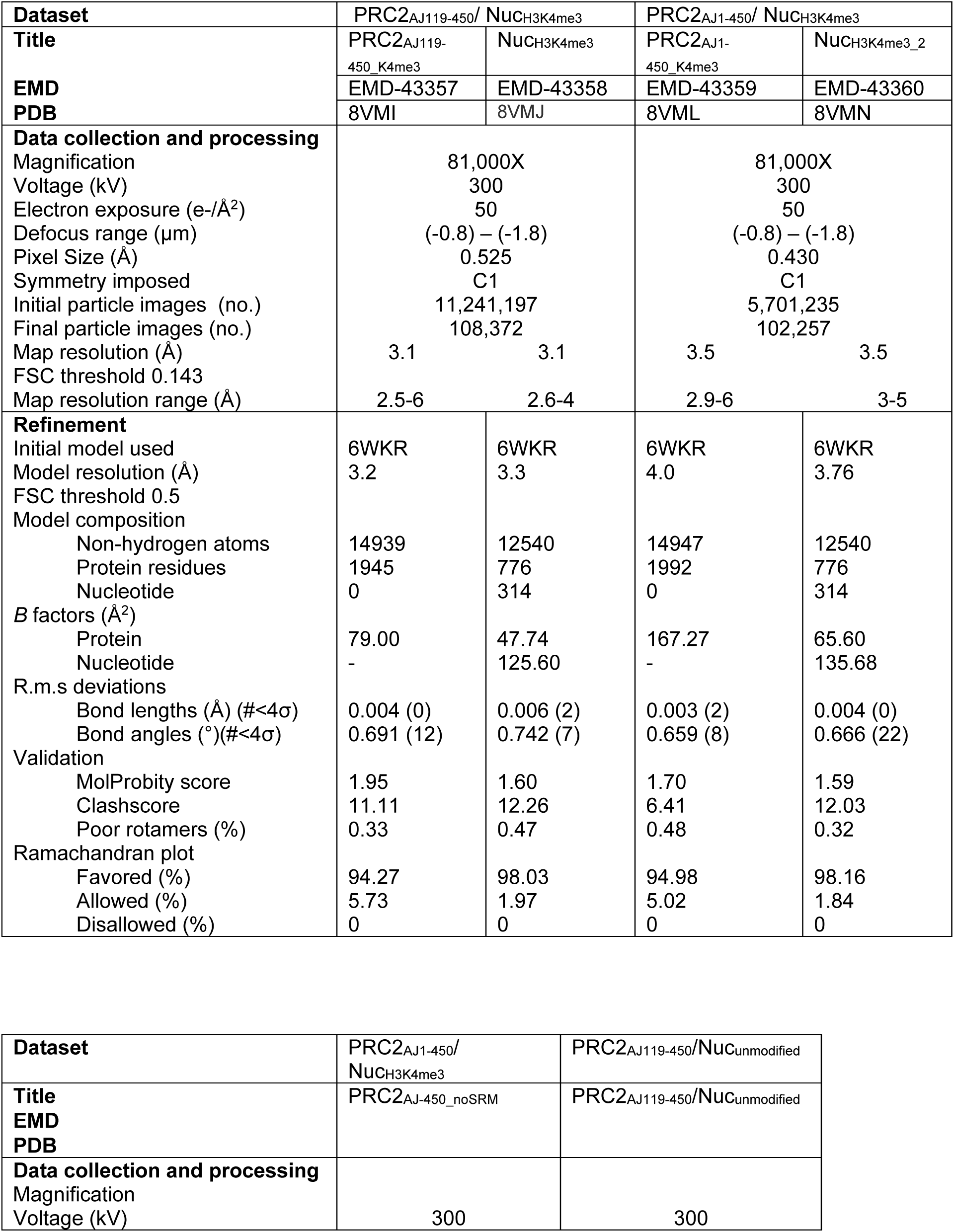
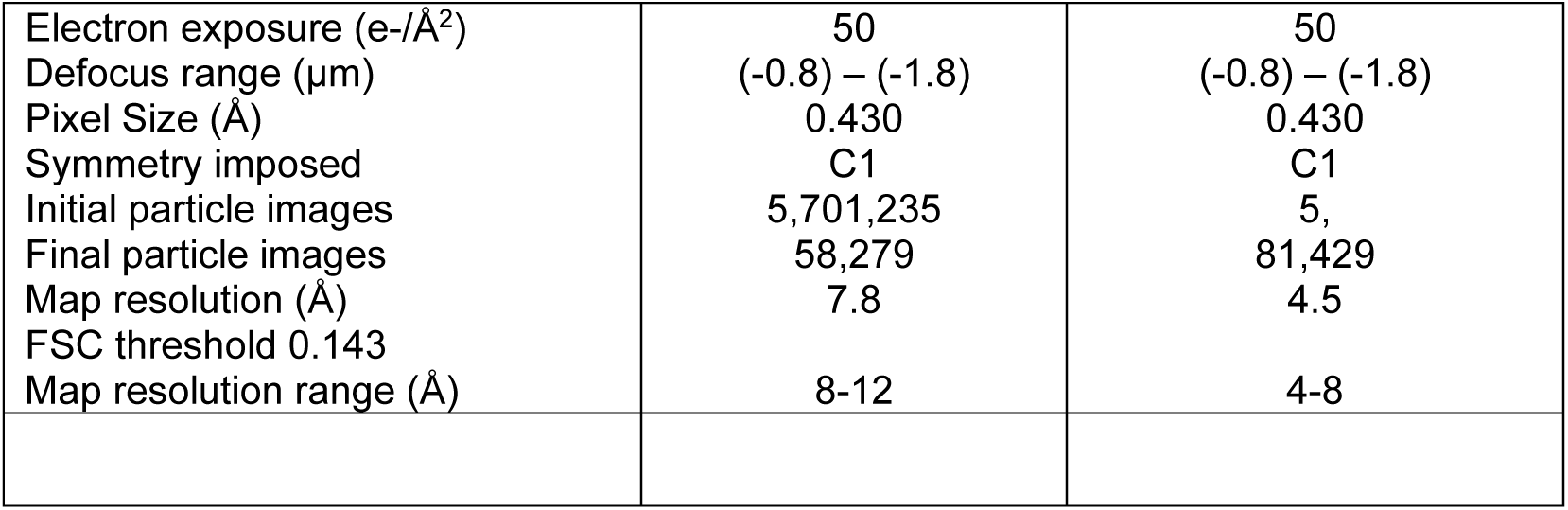
cryo-EM data collection and processing.

## Supplementary Figures

**Figure S1.**
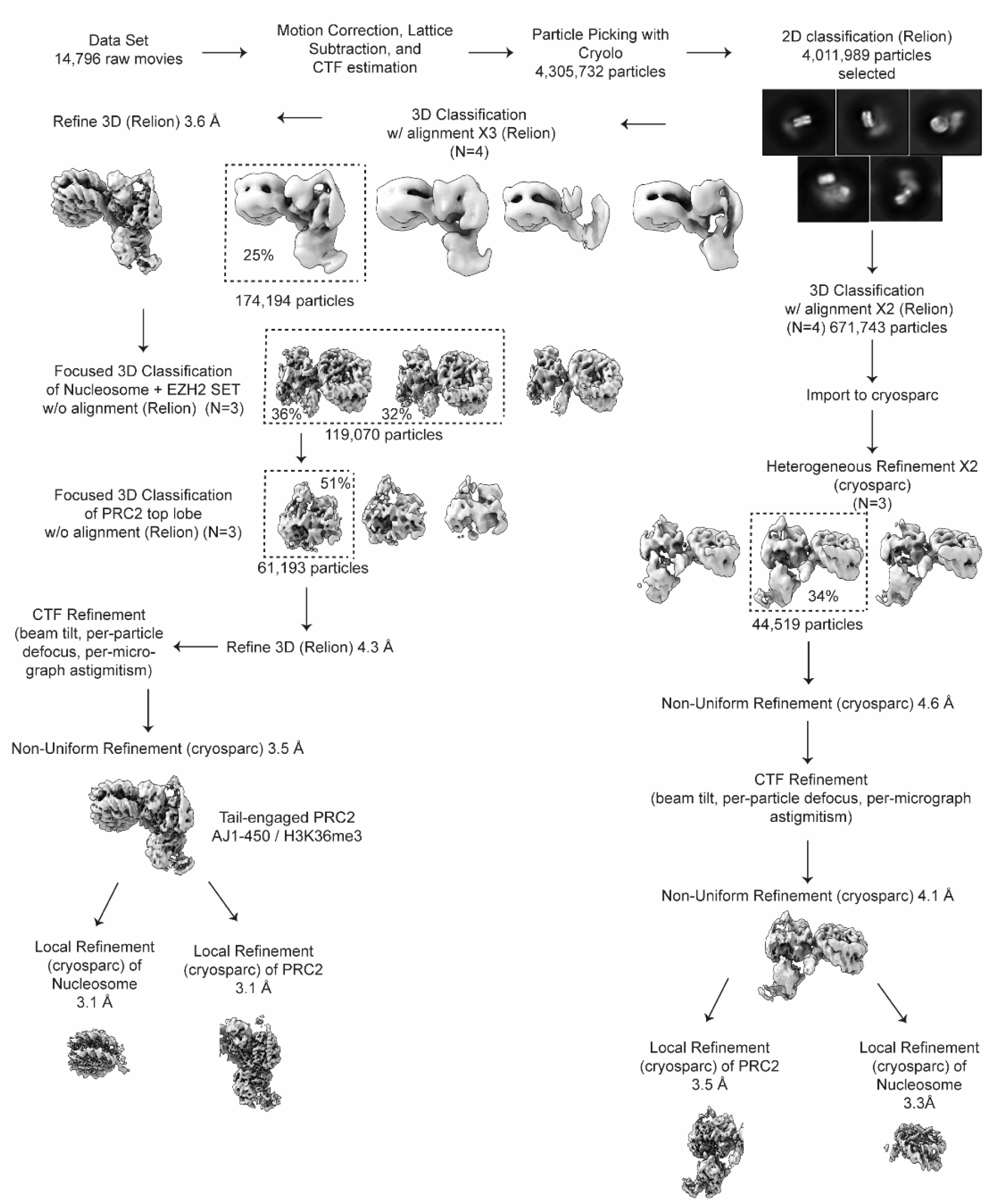
Processing workflow for PRC2_AJ1-450_ bound to H3K36me3 nucleosomes. Data collected for PRC2_AJ1-450_ bound to H3K36me3 nucleosomes was initially processed in RELION. Focused classification around the nucleosome and EZH2 SET domain resulted in two classes, one showing the histone H3 tail engaged and one lacking density for the histone H3 tail. We further refined these particles and performed focused classification around the PRC2 top lobe to obtain a 3.1 Å reconstruction that resolved the histone H3K36me3 side chain (left side of processing diagram). We then used the initial 4,011,989 particles obtained from 2D classification to perform 3D classification, heterogeneous, homogeneous, and local refinements of particles lacking density for the histone H3 tail using Cryo SPARC to resolve the 3.5 Å reconstruction of the “tailless” state (right side of processing diagram).

**Figure S2.**
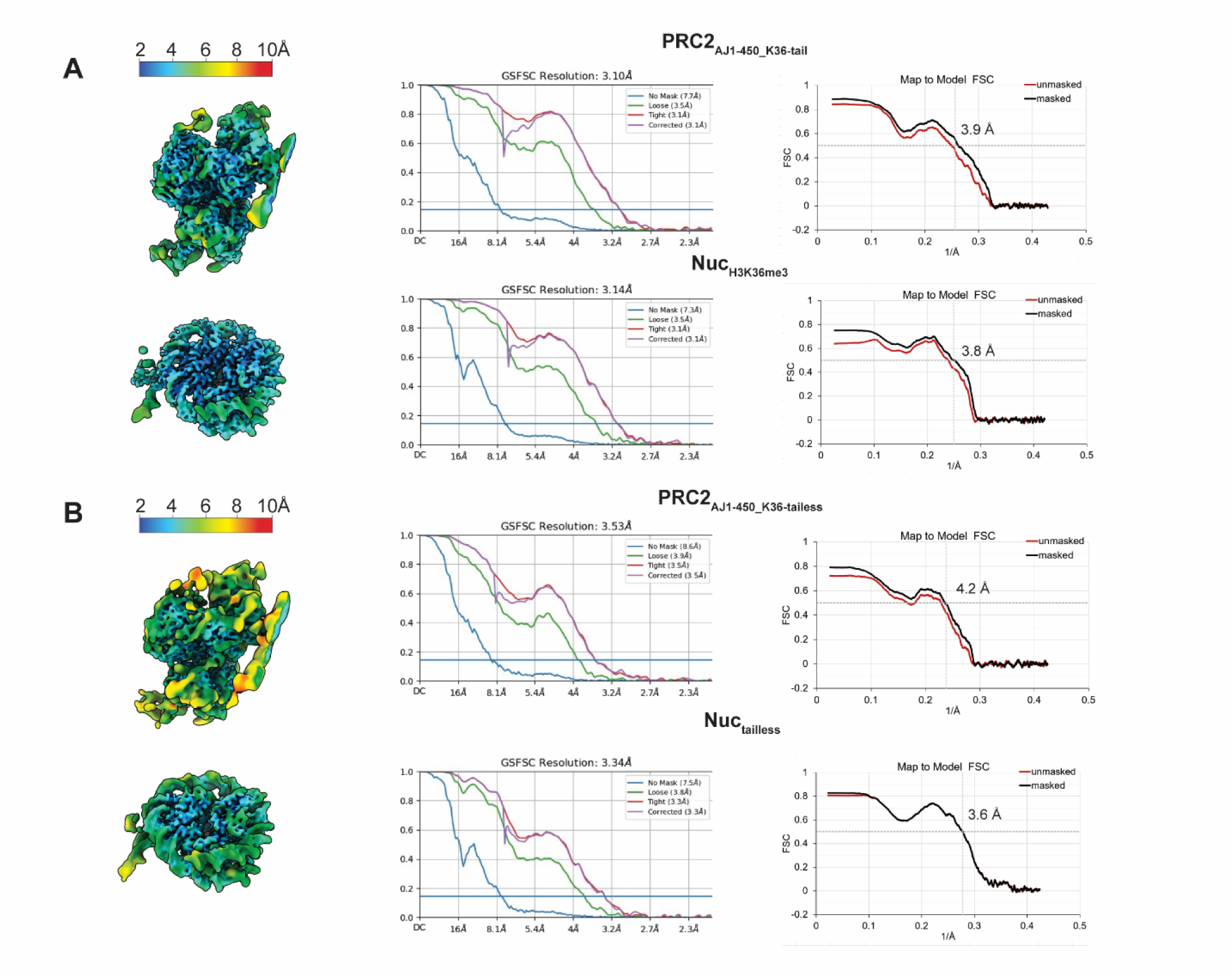
Local resolution mapped onto cryo-EM maps of PRC2_AJ1-450_ bound to H3K36me3-containing nucleosomes obtained from local refinements using masks surrounding PRC2 and nucleosome regions. For each map, the resolution at FSC=0.143 is provided, while the map-to-model FSC plots show the masked resolution at FSC=0.5. Each map has been locally filtered using the local resolution estimate. **A)** PRC2_AJ1-450_ / H3K36me3 “tail-engaged” state **B)** PRC2_AJ1-450_ / H3K36me3 “tailless” state

**Figure S3.**
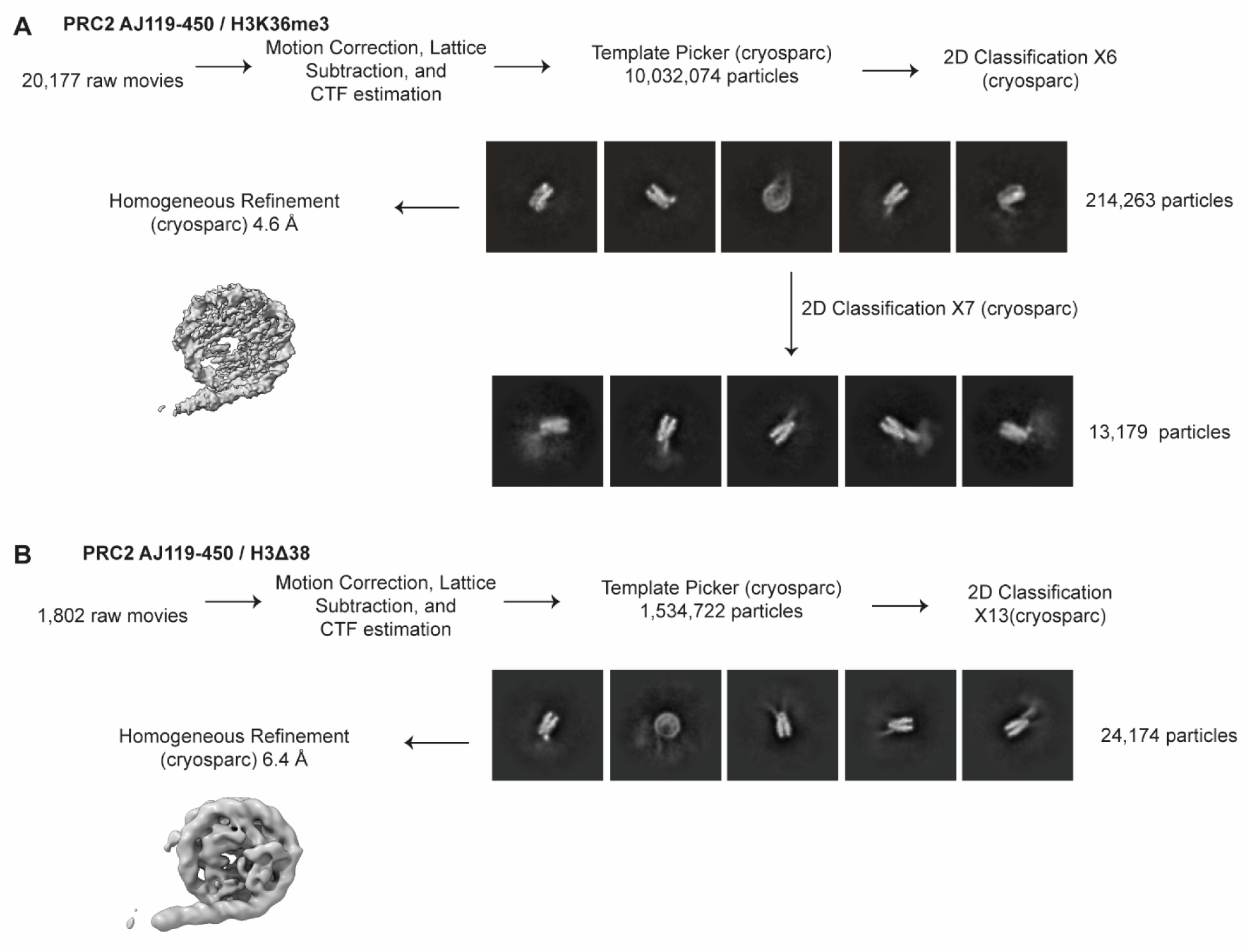
Cryo-EM of PRC2_AJ119-450_ bound to H3K36me3 and to H3Δ38 nucleosomes. **A)** Cryo-EM analysis of PRC2_AJ119-450_ bound to H3K36me3 modified nucleosomes. Data was extensively cleaned by 2D classification trying to bring up density for PRC2. Representative 2D classes after 6 rounds are shown (also in Figure S3A), but the resulting 214,262 particles resulted in a 4.6 Å reconstruction showing only the nucleosome, without additional density for PRC2. Further cleaning of the data by 2D classification (after 13 total rounds) only showed fuzzy density for PRC2. **B)** Cryo-EM of PRC2_AJ119-450_ bound to nucleosomes containing H3Δ38. Data was extensively cleaned by 2D classification trying to bring up density for PRC2. Representative 2D classes after 13 rounds are shown (also in Figure S3B). The resulting 24,172 particles were used to obtain a 6.4 Å reconstruction that shows only the nucleosome, without additional density for PRC2, despite the weak density that can be observed in the 2D classification.

**Figure S4.**
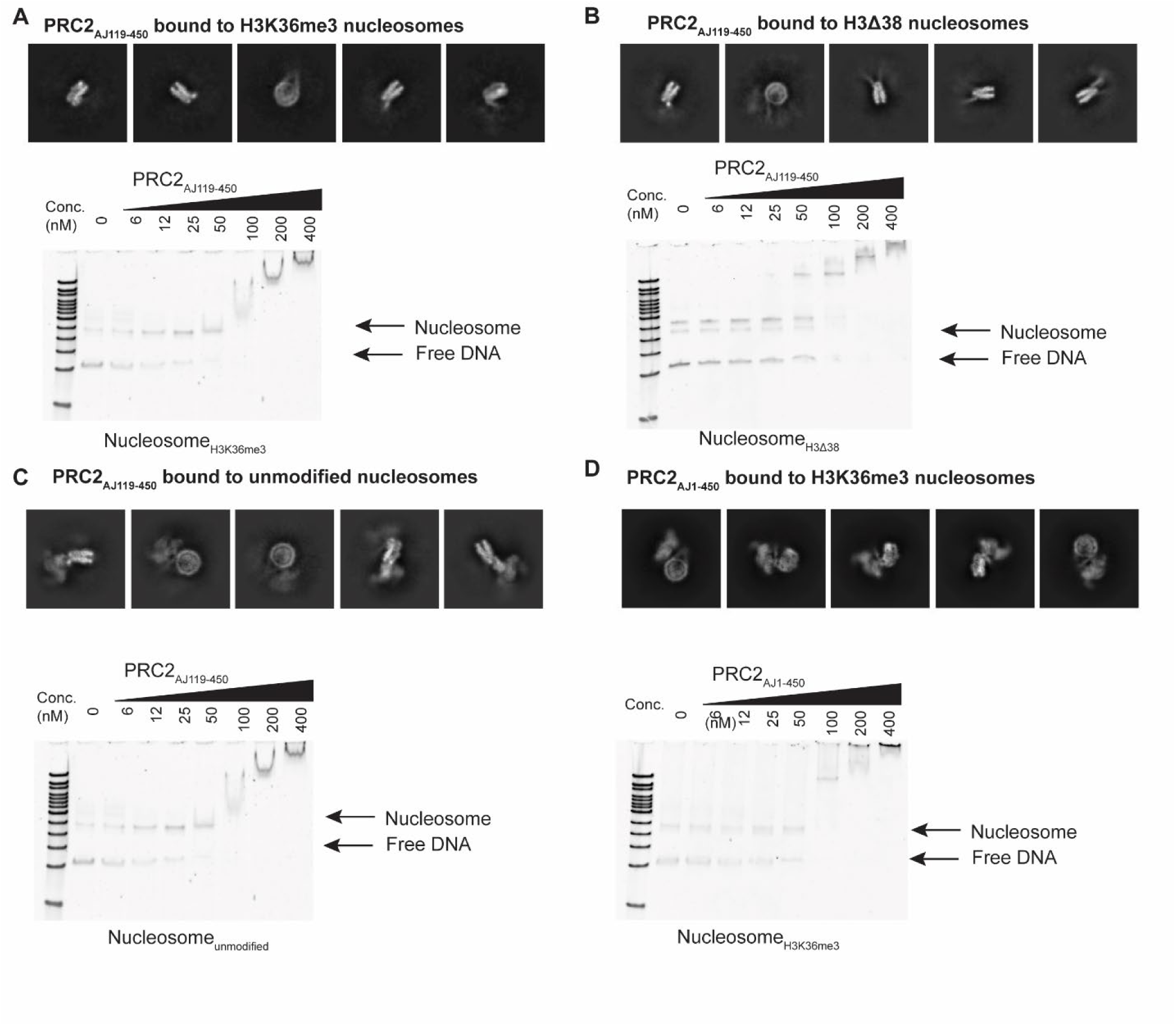
Cryo-EM and nucleosome binding analysis of PRC2 with H3K36me3, unmodified, and H3Δ38 nucleosomes. **A)** Representative 2D classes of PRC2_AJ119-450_ bound to H3K36me3 nucleosomes show absence of strong density for PRC2, despite observed binding in electromobility shift assays. Electromobility shift assays were performed with a titration of 0 to 400 nM PRC2 with 50 nM nucleosome in all cases. **B)** Representative 2D classes of PRC2_AJ119-450_ bound to nucleosomes containing histone H3Δ38 show absence of strong density for PRC2, despite observed binding in electromobility shift assays. **C)** Representative 2D classes of PRC2_AJ119-450_ bound to unmodified nucleosomes that yielded the 4.5 Å reconstruction. **D) Representative 2D classes of** PRC2_AJ1-450_ bound to H3K36me3 nucleosomes that yielded the 3.6 Å tail-engaged reconstruction. In contrast to A and B, C and D show strong density for PRC2. Electromobility shift assays show similar binding affinity for PRC2_AJ119-450_ bound to unmodified nucleosomes or PRC2_AJ1-450_ bound to H3K36me3 nucleosomes to that observed for the PRC2_AJ119-450_/Nucleosome_H3K36me3_ and PRC2_AJ119-450_/Nucleosome_H3Δ38_ assays.

**Figure S5.**
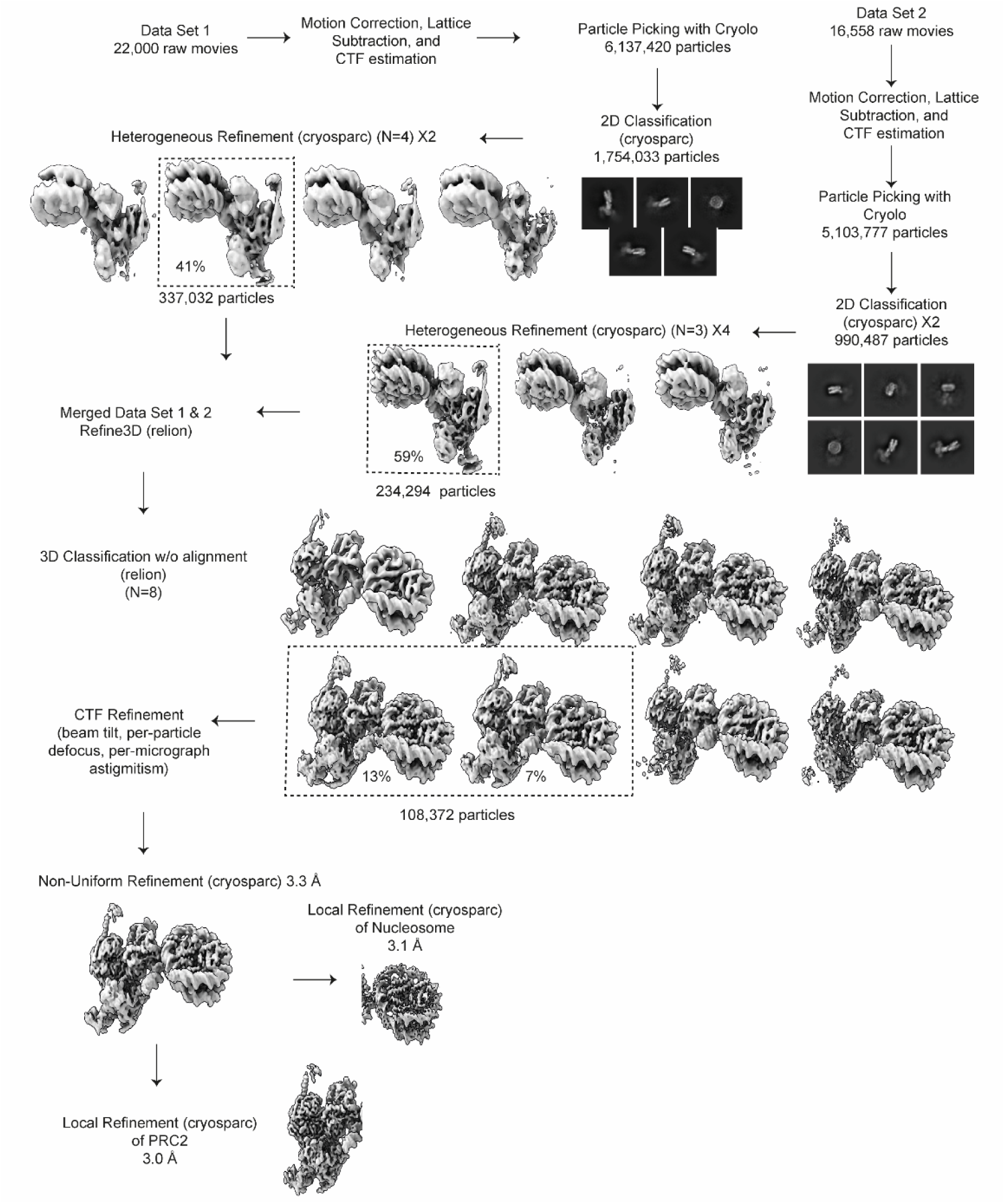
Processing workflow for PRC2_AJ119-450_ bound to H3K4me3 nucleosomes. Data collected for PRC2_AJ119-450_ bound to H3K4me3 nucleosomes was initially processed in cryosparc. Two data sets were merged after 2D classification and one round of heterogeneous refinement. Particles were imported into RELION for 3D Refinement followed by 3D classification without alignment. All 3D classes show density for the histone H3 tail and absence of density for the EZH2 SRM. Classes were selected showing the strongest density for H3K4me3 bound to EED. After CTF Refinement, particles were imported into cryosparc for non-uniform refinement and local refinements around the nucleosome and PRC2 regions.

**Figure S6.**
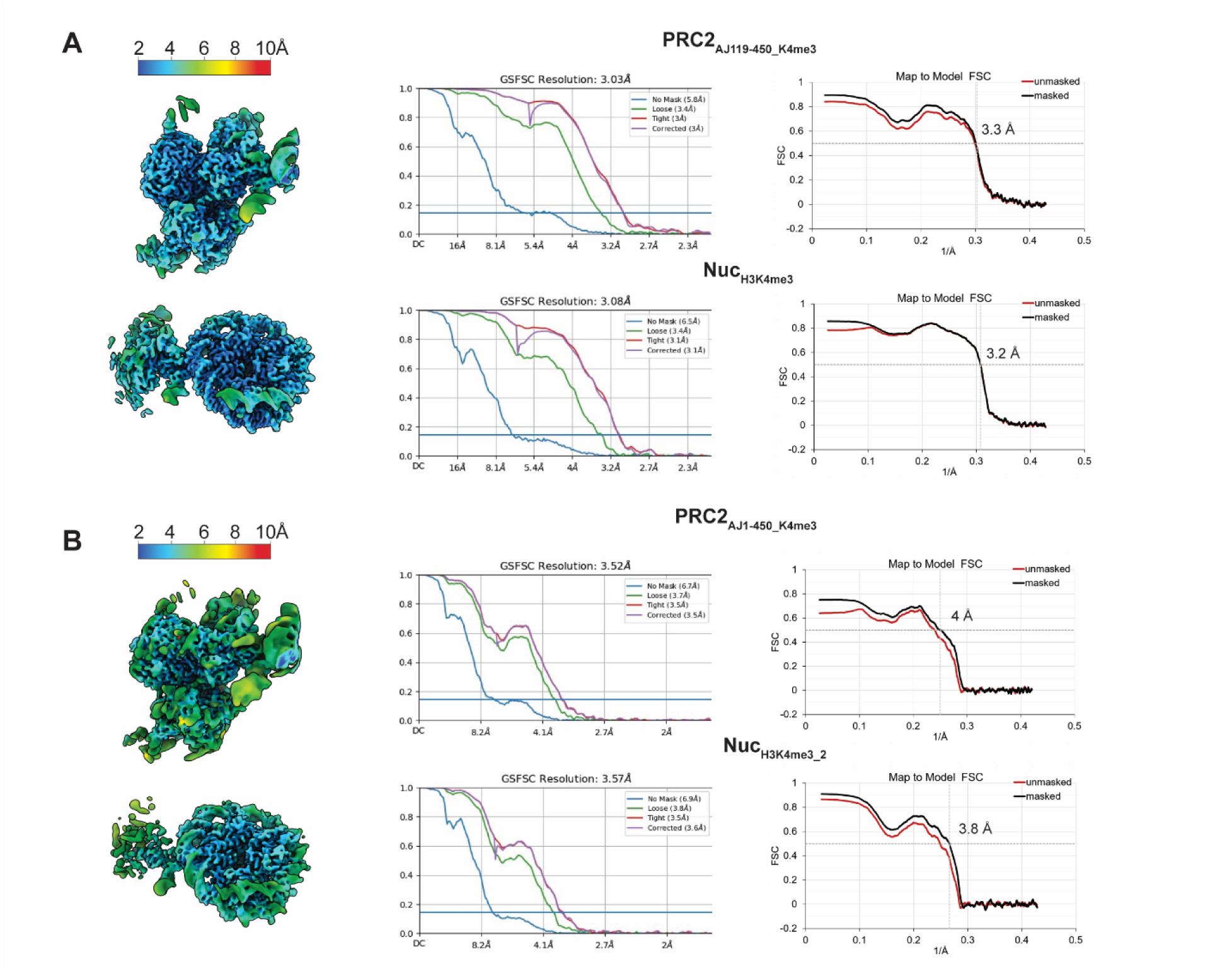
Local resolution mapped onto cryo-EM maps of PRC2 complexes bound to H3K4me3-containing nucleosomes obtained from local refinements using masks surrounding PRC2 or nucleosome regions. For each map, the resolution at FSC=0.143 is provided, while the map-to-model FSC plots show the masked resolution at FSC=0.5. Each map has been locally filtered using the local resolution estimate. **A)** PRC2_AJ119-450_ / H3K4me3 **B)** PRC2_AJ1-450_ / H3K4me3

**Figure S7.**
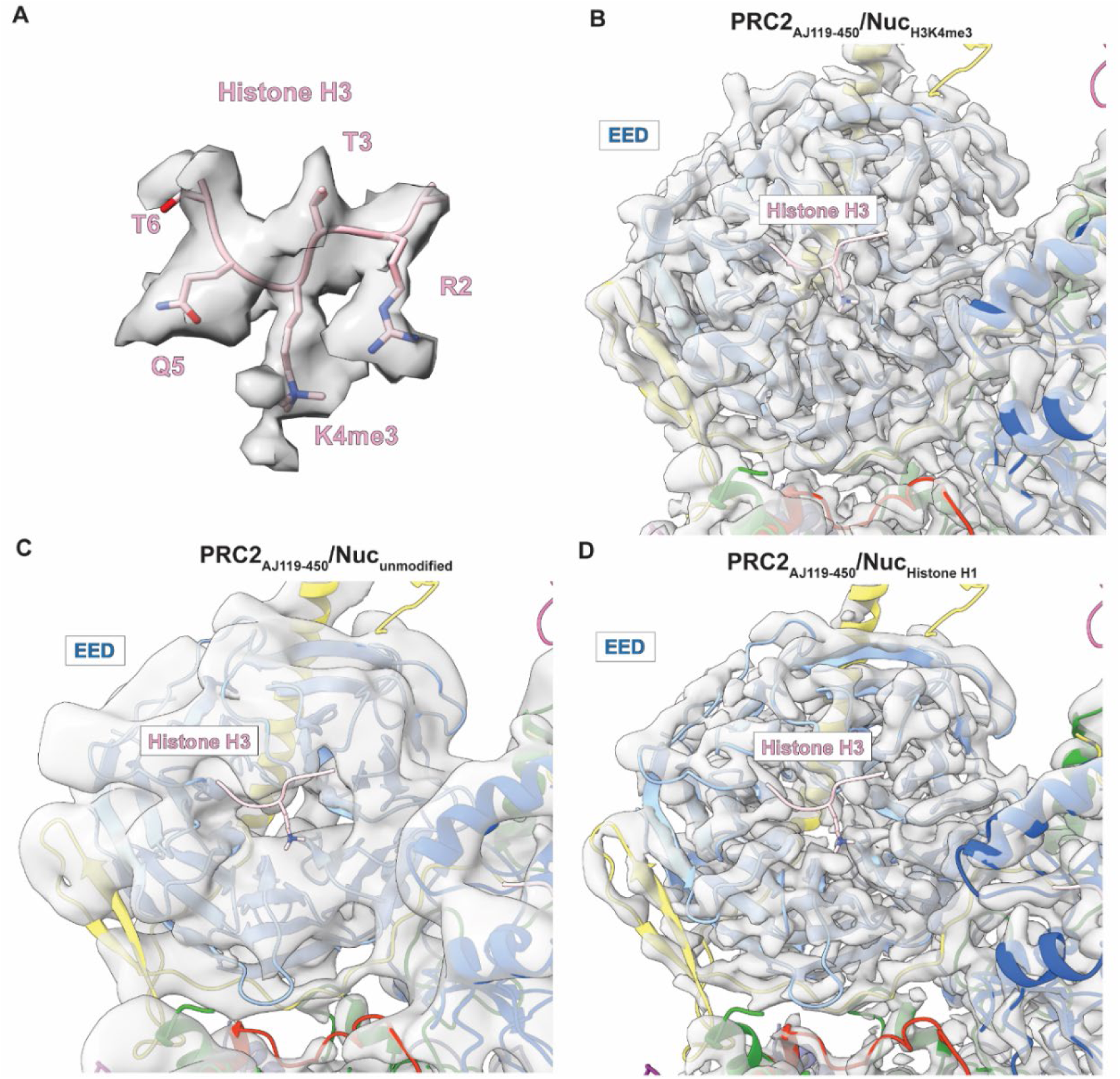
Comparison of cryo-EM density for the EED allosteric site in structures of PRC2_AJ119-450_ bound to H3K4me3 or to unmodified nucleosomes. **A)** Density surrounding the H3K4me3 bound to the EED for the PRC2_AJ119-450_ bound to a H3K4me3-modified nucleosome (resolution ∼ 3.1 Å) shown at threshold of 0.232 **B)** Cryo-EM density map (transparent grey) and model of the EED allosteric site and the bound methylated peptide for PRC2_AJ119-450_ bound to a H3K4me3-modified nucleosome (resolution ∼ 3.1 Å). The model is shown in ribbon representation colored by subunit, withEED shown in light blue and the histone H3K4me3 peptide shown in pink. **C)** Corresponding view of a reconstruction obtained for PRC2_AJ119-450_ bound to a unmodified nucleosome (resolution ∼ 4.5 Å) showing lack of density in the EED allosteric pocket . **D)** Corresponding view of our recently reported reconstruction obtained for PRC2_AJ119-450_ bound to a an unmodified nucleosome that additionally contained the linker histone H1 (resolution ∼ 3.6 Å), also showing lack of density in the EED allosteric pocket.

**Figure S8.**
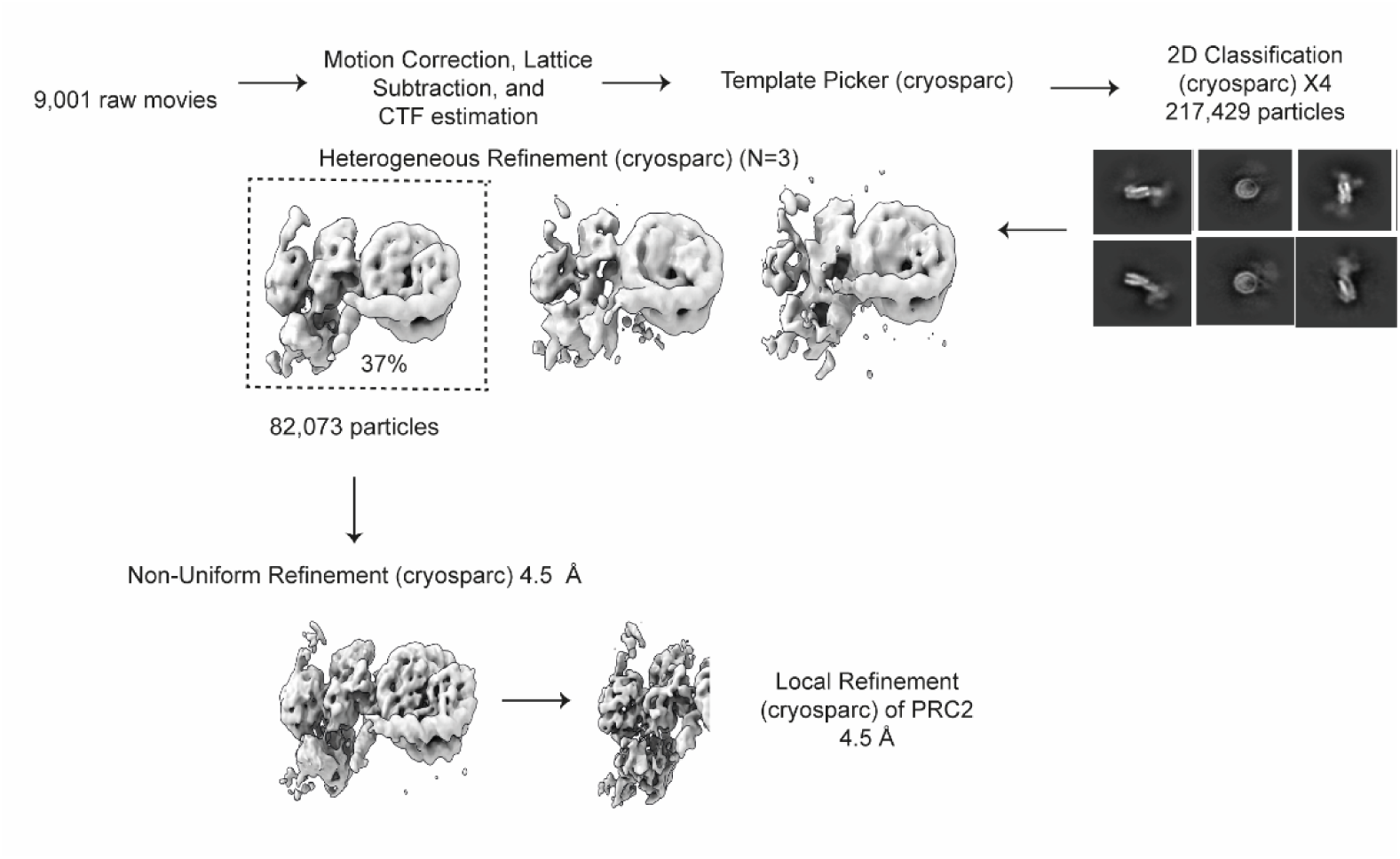
Processing workflow for PRC2_AJ119-450_ bound to unmodified nucleosomes. Data collected for PRC2_AJ119-450_ bound to unmodified nucleosomes was processed in cryosparc following standard workflow. Particles were subjected to 2D classification, followed by heterogeneous refinement to remove damaged complexes. Non-uniform refinement was performed followed by local refinement around PRC2.

**Figure S9.**
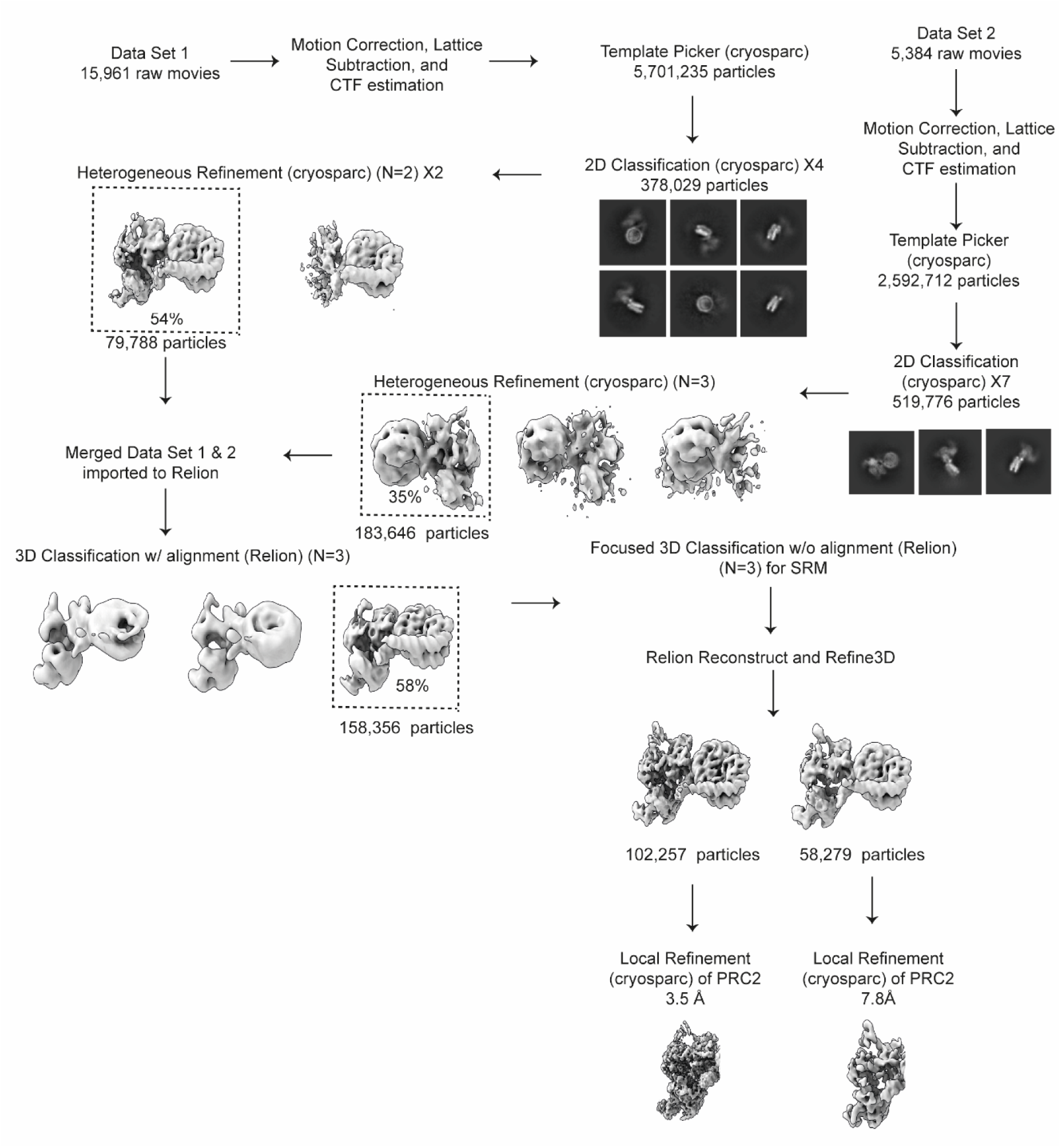
Processing workflow for PRC2_AJ1-450_ bound to H3K4me3 nucleosomes. Data collected for PRC2_AJ1-450_ bound to H3K36me3 nucleosomes was processed in cryosparc. Two data sets were merged after 2D classification and heterogeneous refinement. Particles were imported into RELION and subjected to 3D classification without alignment. Focused classification was performed using a mask around the allosteric site and EZH2 SRM regions resulting in two major classes showing the presence of absence of the SRM. Each class was refined separately and imported into cryosparc for local refinement around the PRC2 region.

**Figure S10.**
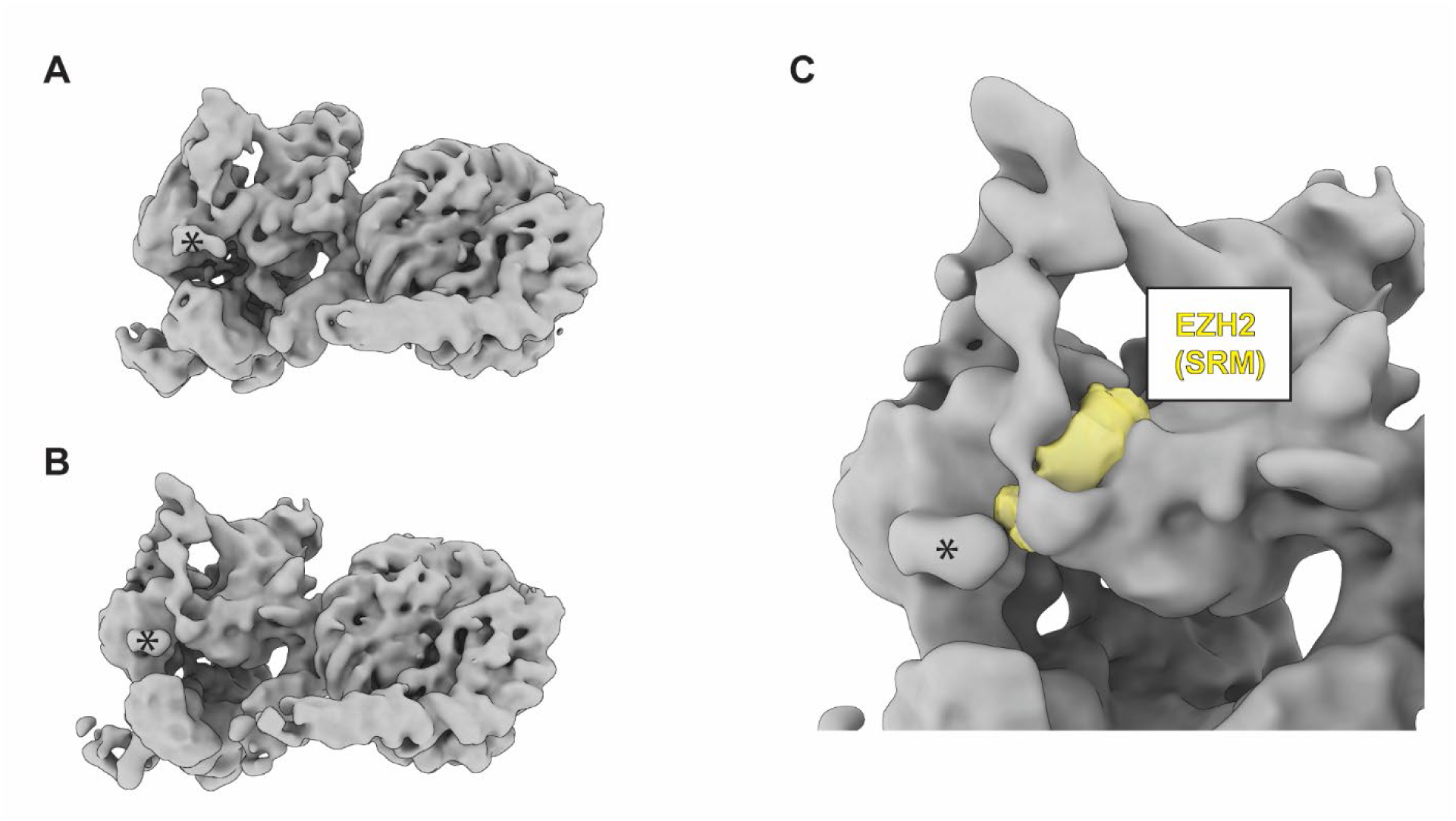
PRC2_AJ1-450_ bound to H3K4me3 nucleosomes with unfolded SRM. **A)** “Activated” PRC2_AJ1-450_ / H3K4me3 cryo-EM map lowpass filtered to 8 Å. **B)** PRC2_AJ1-450_ / H3K4me3 cryo-EM map, which lacks density for the SRM, lowpass filtered to 8 Å. **C)** Close up view of the allosteric site of the PRC2_AJ1-450_ / H3K4me3 cryo-EM map (grey) showing the corresponding SRM density (yellow) in the “activated” PRC2_AJ1-450_ / H3K4me3 cryo-EM map. Both maps contain an unassigned density marked with asterisk that is not apparent in any of the other cryo-EM density maps of PRC2/nucleosome complexes.

## References

1. Cao, R. et al. Role of Histone H3 Lysine 27 Methylation in Polycomb-Group Silencing. Science 298, 1039–1043 (2002).

2. Hauri, S. et al. A High-Density Map for Navigating the Human Polycomb Complexome. Cell Rep 17, 583–595 (2016).

3. Grijzenhout, A. et al. Functional analysis of AEBP2, a PRC2 Polycomb protein, reveals a Trithorax phenotype in embryonic development and in ESCs. Development 143, 2716–2723 (2016).

4. Poepsel, S., Kasinath, V. & Nogales, E. Cryo-EM structures of PRC2 simultaneously engaged with two functionally distinct nucleosomes. Nature Structural & Molecular Biology 25, 154–162 (2018).

5. Kasinath, V. et al. JARID2 and AEBP2 regulate PRC2 in the presence of H2AK119ub1 and other histone modifications. Science 371, eabc3393 (2021).

6. Kasinath, V. et al. Structures of human PRC2 with its cofactors AEBP2 and JARID2. Science 359, 940–944 (2018).

7. Jiao, L. & Liu, X. Structural basis of histone H3K27 trimethylation by an active polycomb repressive complex 2. Science 350, aac4383 (2015).

8. Oksuz, O. et al. Capturing the onset of PRC2-mediated repressive domain formation. Molecular cell 70, 1149–1162. e5 (2018).

9. Margueron, R. et al. Role of the polycomb protein EED in the propagation of repressive histone marks. Nature 461, 762–767 (2009).

10. Yuan, W. et al. Dense Chromatin Activates Polycomb Repressive Complex 2 to Regulate H3 Lysine 27 Methylation. Science 337, 971–975 (2012).

11. Justin, N. et al. Structural basis of oncogenic histone H3K27M inhibition of human polycomb repressive complex 2. Nature Communications 7, 11316 (2016).

12. Lee, C.-H. et al. Allosteric activation dictates PRC2 activity independent of its recruitment to chromatin. Molecular cell 70, 422–434. e6 (2018).

13. Sanulli, S. et al. Jarid2 Methylation via the PRC2 Complex Regulates H3K27me3 Deposition during Cell Differentiation. Molecular Cell 57, 769–783 (2015).

14. Zhang, Q. et al. PALI1 facilitates DNA and nucleosome binding by PRC2 and triggers an allosteric activation of catalysis. Nature Communications 12, 4592 (2021).

15. Cooper, S. et al. Jarid2 binds mono-ubiquitylated H2A lysine 119 to mediate crosstalk between Polycomb complexes PRC1 and PRC2. Nature Communications 7, 13661 (2016).

16. Kalb, R. et al. Histone H2A monoubiquitination promotes histone H3 methylation in Polycomb repression. Nature Structural & Molecular Biology 21, 569–571 (2014).

17. Sauer, P.V. et al. Activation of automethylated PRC2 by dimerization on chromatin. bioRxiv, 2023.10.12.562141 (2023).

18. Finogenova, K. et al. Structural basis for PRC2 decoding of active histone methylation marks H3K36me2/3. eLife 9, e61964 (2020).

19. Grau, D. et al. Structures of monomeric and dimeric PRC2: EZH1 reveal flexible modules involved in chromatin compaction. Nature communications 12, 714 (2021).

20. Song, J. et al. Structural basis for inactivation of PRC2 by G-quadruplex RNA. Science 381, 1331–1337 (2023).

21. Kaneko, S., Son, J., Bonasio, R., Shen, S.S. & Reinberg, D. Nascent RNA interaction keeps PRC2 activity poised and in check. Genes & development 28, 1983–1988 (2014).

22. Beltran, M. et al. The interaction of PRC2 with RNA or chromatin is mutually antagonistic. Genome research 26, 896–907 (2016).

23. Wang, X. et al. Molecular analysis of PRC2 recruitment to DNA in chromatin and its inhibition by RNA. Nature Structural & Molecular Biology 24, 1028–1038 (2017).

24. Yan, J., Dutta, B., Hee, Y.T. & Chng, W.-J. Towards understanding of PRC2 binding to RNA. RNA biology 16, 176–184 (2019).

25. Schmitges, F.W. et al. Histone methylation by PRC2 is inhibited by active chromatin marks. Molecular cell 42, 330–341 (2011).

26. Yuan, W. et al. H3K36 Methylation Antagonizes PRC2-mediated H3K27 Methylation Journal of Biological Chemistry 286, 7983–7989 (2011).

27. Schneider, R. et al. Histone H3 lysine 4 methylation patterns in higher eukaryotic genes. Nature Cell Biology 6, 73–77 (2004).

28. Pokholok, D.K. et al. Genome-wide Map of Nucleosome Acetylation and Methylation in Yeast. Cell 122, 517–527 (2005).

29. Bernstein, B.E. et al. Genomic Maps and Comparative Analysis of Histone Modifications in Human and Mouse. Cell 120, 169–181 (2005).

30. Vakoc, C.R., Sachdeva, M.M., Wang, H. & Blobel, G.A. Profile of Histone Lysine Methylation across Transcribed Mammalian Chromatin. Molecular and Cellular Biology 26, 9185–9195 (2006).

31. Santos-Rosa, H. et al. Active genes are tri-methylated at K4 of histone H3. Nature 419, 407–11 (2002).

32. Kizer, K.O. et al. A Novel Domain in Set2 Mediates RNA Polymerase II Interaction and Couples Histone H3 K36 Methylation with Transcript Elongation. Molecular and Cellular Biology 25, 3305–3316 (2005).

33. Bannister, A.J. et al. Spatial distribution of di- and tri-methyl lysine 36 of histone H3 at active genes. J Biol Chem 280, 17732–6 (2005).

34. Ernst, J. & Kellis, M. Discovery and characterization of chromatin states for systematic annotation of the human genome. Nature Biotechnology 28, 817–825 (2010).

35. Kharchenko, P.V. et al. Comprehensive analysis of the chromatin landscape in Drosophila melanogaster. Nature 471, 480–485 (2011).

36. Gaydos, Laura J., Rechtsteiner, A., Egelhofer, Thea A., Carroll, Coleen R. & Strome, S. Antagonism between MES-4 and Polycomb Repressive Complex 2 Promotes Appropriate Gene Expression in C. elegans Germ Cells. Cell Reports 2, 1169–1177 (2012).

37. Musselman, C.A. et al. Molecular basis for H3K36me3 recognition by the Tudor domain of PHF1. Nature Structural & Molecular Biology 19, 1266–1272 (2012).

38. Venkatesh, S. & Workman, J.L. Set2 mediated H3 lysine 36 methylation: regulation of transcription elongation and implications in organismal development. WIREs Developmental Biology 2, 685–700 (2013).

39. Li, J., Moazed, D. & Gygi, S.P. Association of the Histone Methyltransferase Set2 with RNA Polymerase II Plays a Role in Transcription Elongation *. Journal of Biological Chemistry 277, 49383–49388 (2002).

40. Han, B.G. et al. Long shelf-life streptavidin support-films suitable for electron microscopy of biological macromolecules. J Struct Biol 195, 238–244 (2016).

41. Au - Cookis, T., et al. Streptavidin-Affinity Grid Fabrication for Cryo-Electron Microscopy Sample Preparation. JoVE, e66197 (2023).

42. Choi, J. et al. DNA binding by PHF1 prolongs PRC2 residence time on chromatin and thereby promotes H3K27 methylation. Nature Structural & Molecular Biology 24, 1039–1047 (2017).

43. Youmans, D.T., Schmidt, J.C. & Cech, T.R. Live-cell imaging reveals the dynamics of PRC2 and recruitment to chromatin by SUZ12-associated subunits. Genes Dev 32, 794–805 (2018).

44. Højfeldt, J.W. et al. Accurate H3K27 methylation can be established de novo by SUZ12-directed PRC2. Nat Struct Mol Biol 25, 225–232 (2018).

45. Wang, H. et al. H3K4me3 regulates RNA polymerase II promoter-proximal pause-release. Nature 615, 339–348 (2023).

46. Hughes, A.L., Kelley, J.R. & Klose, R.J. Understanding the interplay between CpG island-associated gene promoters and H3K4 methylation. Biochimica et Biophysica Acta (BBA) - Gene Regulatory Mechanisms 1863, 194567 (2020).

47. Vermeulen, M. et al. Selective Anchoring of TFIID to Nucleosomes by Trimethylation of Histone H3 Lysine 4. Cell 131, 58–69 (2007).

48. Lauberth, Shannon M. et al. H3K4me3 Interactions with TAF3 Regulate Preinitiation Complex Assembly and Selective Gene Activation. Cell 152, 1021–1036 (2013).

49. Xu, C. et al. Binding of different histone marks differentially regulates the activity and specificity of polycomb repressive complex 2 (PRC2). Proceedings of the National Academy of Sciences 107, 19266–19271 (2010).

50. Ferrari, K.J. et al. Polycomb-dependent H3K27me1 and H3K27me2 regulate active transcription and enhancer fidelity. Mol Cell 53, 49–62 (2014).

51. Streubel, G. et al. The H3K36me2 methyltransferase Nsd1 demarcates PRC2-mediated H3K27me2 and H3K27me3 domains in embryonic stem cells. Molecular cell 70, 371–379. e5 (2018).

52. Sneeringer, C.J. et al. Coordinated activities of wild-type plus mutant EZH2 drive tumor-associated hypertrimethylation of lysine 27 on histone H3 (H3K27) in human B-cell lymphomas. Proc Natl Acad Sci U S A 107, 20980–5 (2010).

53. Conway, E. et al. A Family of Vertebrate-Specific Polycombs Encoded by the LCOR/LCORL Genes Balance PRC2 Subtype Activities. Mol Cell 70, 408–421.e8 (2018).

54. Steiner, L.A., Schulz, V.P., Maksimova, Y., Wong, C. & Gallagher, P.G. Patterns of histone H3 lysine 27 monomethylation and erythroid cell type-specific gene expression. J Biol Chem 286, 39457–65 (2011).

55. Pasini, D. et al. JARID2 regulates binding of the Polycomb repressive complex 2 to target genes in ES cells. Nature 464, 306–10 (2010).

56. Li, G. et al. Jarid2 and PRC2, partners in regulating gene expression. Genes Dev 24, 368–80 (2010).

57. Zheng, S.Q. et al. MotionCor2: anisotropic correction of beam-induced motion for improved cryo-electron microscopy. Nature Methods 14, 331–332 (2017).

58. Rohou, A. & Grigorieff, N. CTFFIND4: Fast and accurate defocus estimation from electron micrographs. J Struct Biol 192, 216–21 (2015).

59. Wagner, T. et al. SPHIRE-crYOLO is a fast and accurate fully automated particle picker for cryo-EM. Communications Biology 2, 218 (2019).

60. Scheres, S.H.W. RELION: Implementation of a Bayesian approach to cryo-EM structure determination. Journal of Structural Biology 180, 519–530 (2012).

61. Croll, T. ISOLDE: a physically realistic environment for model building into low-resolution electron-density maps. Acta Crystallographica Section D 74, 519–530 (2018).

62. Pettersen, E.F. et al. UCSF ChimeraX: Structure visualization for researchers, educators, and developers. Protein Sci 30, 70–82 (2021).

63. Emsley, P. & Cowtan, K. Coot: model-building tools for molecular graphics. Acta Crystallogr D Biol Crystallogr 60, 2126–32 (2004).

64. Adams, P.D. et al. PHENIX: building new software for automated crystallographic structure determination. Acta Crystallogr D Biol Crystallogr 58, 1948–54 (2002).

